# Genomic ancestry, diet and microbiomes of Upper Palaeolithic hunter-gatherers from San Teodoro cave (Sicily, Italy)

**DOI:** 10.1101/2021.12.08.471745

**Authors:** Gabriele Scorrano, Sofie Holtsmark Nielsen, Domenico Lo Vetro, Meaghan Mackie, Ashot Margaryan, Anna K. Fotakis, Cristina Martínez-Labarga, Pier Francesco Fabbri, Morten E. Allentoft, Marialetizia Carra, Fabio Martini, Olga Rickards, Jesper V. Olsen, Enrico Cappellini, Martin Sikora

**Affiliations:** Section for Evolutionary Genomics, Globe Institute, University of Copenhagen, Copenhagen, Denmark; Lundbeck Foundation GeoGenetics Centre, Globe Institute, University of Copenhagen, Copenhagen, Denmark; Dipartimento di Storia, Archeologia, Geografia, Arte e Spettacolo (SAGAS), University of Florence, Florence, Italy; Museo e Istituto Fiorentino di Preistoria, Florence, Italy; Novo Nordisk Foundation Center for Protein Research, Faculty of Health and Medical Sciences, University of Copenhagen, Copenhagen, Denmark; Center for Evolutionary Hologenomics, University of Copenhagen, Copenhagen, Denmark; Centro di Antropologia Molecolare per lo studio del DNA antico, Dipartimento di Biologia, University of Rome Tor Vergata, Rome, Italy; Dipartimento di Beni Culturali, University of Salento, Lecce, Italy; Trace and Environmental DNA (TrEnD) Laboratory, School of Molecular and Life Sciences, Curtin University, Perth, Australia; DANTE: Diet and Ancient Technology Laboratory, Sapienza University of Rome, Rome, Italy

**Keywords:** Upper Palaeolithic, human aDNA, metagenomics, palaeoproteomics, multi-omics approach

## Abstract

Recent improvements in the analysis of ancient biomolecules from human remains and associated dental calculus have provided new insights into the prehistoric diet and past genetic diversity of our species. Here we present a “multi-omics” study, integrating genomic and proteomic analyses of two post-Last Glacial Maximum (LGM) individuals from San Teodoro cave (Italy), to reconstruct their lifestyle and the post-LGM resettlement of Europe. Our analyses show genetic homogeneity in Sicily during the Palaeolithic, representing a hitherto unknown Italian genetic lineage within the previously identified “Villabruna cluster”. We argue that this lineage took refuge in Italy during the LGM, followed by a subsequent spread to central-western Europe. Multi-omics analysis of dental calculus showed a diet rich of animal proteins which is also reflected on the oral microbiome composition. Our results demonstrate the power of using a multi-omics approach in the study of prehistoric human populations.

## Introduction

In recent years, improvement of ancient DNA (aDNA) methods have given unprecedented insights into the past population dynamics of our species^1,2^. To date, most aDNA research focus on the dispersal of anatomically modern humans across the world, exploring the history of migration routes and admixture events which shaped extant human genetic variability. Recent studies have clarified the peopling of Western Eurasia during the Upper Palaeolithic after modern humans migrated out of Africa^1^. While the earliest peoples arriving in Europe around 45 kya appear not to have contributed ancestry to later groups^2^; the genome of an early European individual from Kostenki 14, dated to around 37 kya, demonstrated that the ancestral European gene pool was already established by that time^3^.

Analyses of genomic data from a larger set of pre-Neolithic Europeans documented complex population structure among early Europeans, involving multiple deeply diverged lineages. Among the identified pre- and post-LGM genetic clusters, two lineages, El Mirón and Villabruna^3^, share the highest number of alleles with present-day Europeans. Both lineages were widespread in Europe during the major warming period after the LGM, the Bølling-Allerød interstadial^4^. El Mirón (in Europe around 19 to 14 kya) is associated with the postglacial spread of Magdalenian culture from southwestern European refuges and shares genetic ancestry with the pre-LGM Goyet-Q116-1^3^. The Villabruna genetic cluster (14 to 7 kya) was found to be the predominant ancestry cluster among Western and Central European hunter-gatherers and is associated with the Azilian, Epipaleolithic, Epigravettian and Mesolithic cultures in Europe^3^. Nevertheless, a more complete picture of the post-LGM population history of western Eurasia remains elusive, as fossils from Southern Europe are still underrepresented in genomic studies.

Meanwhile, recent investigations have shown that dental calculus, a complex and calcified bacterial biofilm formed from dental plaque, saliva and gingival crevicular fluid, is rich in aDNA and proteins^5^. Accordingly, calculus is particularly valuable for characterizing diet, oral microbiome and oral disease in ancient populations^5^. Diet is one of the most important lifestyle factors determining human health, playing a pivotal role in shaping the composition of oral microbiomes. Changes in the human diet have implications on the evolution and ecology of the oral microbiome^6^, which in turn can affect gene expression in the immune response system^7^. Ultimately, a deeper knowledge of diet and nutrition becomes necessary to solve the complex co-evolution of oral microbiomes and their human hosts.

Dietary information has so far mostly been obtained by stable isotope analysis, which cannot identify the animal and plant species used as diet resources^8,9^. To overcome this limitation and better characterize the ancient diet and oral microbiomes, dental calculus should be analysed by a double approach to achieve palaeogenomic and palaeoproteomic profiling. These methods are complementary in the identification of species characterizing the oral microbiome and those consumed for nutrition. Furthermore, the proteomic analysis allows for the reconstruction of both the pathogenic action and the immune response ongoing between an oral microbe and its host, based on the detection of process-specific protein profiles^10^. The combined analysis of different ancient biomolecules can therefore provide a wider perspective and answer complex biological questions about past humans^11^.

Here, we combine analyses of human aDNA from petrous bones, with microbial aDNA and food source ancient proteins from dental calculus (Figure S1), isolated from two Late Upper Palaeolithic (Late Epigravettian) hunter-gatherer individuals, dated to 15,322 – 14,432 cal. BP (SI Appendix), from San Teodoro cave (Sicily, Italy; Figure S2, SI Appendix). We used these datasets to investigate genetic ancestry and dispersal of southern European hunter-gatherers after the LGM, and to reconstruct their oral microbiomes and dietary lifestyle. Given the complexity of its archaeological records, Southern Italy is one of the key geographic areas for understanding human and biological responses to postglacial climate evolution in Europe.

## Results

### Human DNA

#### Data generation and authentication

We extracted DNA from the petrous bones of two Palaeolithic individuals from San Teodoro (Sicily, Italy), and sequenced the obtained libraries to 0.5X (San Teodoro 3 – ST3) and 0.1X (San Teodoro 5 – ST5) average genomic coverage using shotgun sequencing (Table S5). Both samples showed deamination pattern and read lengths typically found in ancient remains (Figure S7a), and low rates of contamination with modern human DNA (ST3: 0.4 % mitochondrial DNA (mtDNA) and 2.3% X chromosome; ST5: 1.4% mtDNA; Table S5, Figure S7b), supporting the authenticity of the generated data.

#### Molecular sex, kinship analysis and uniparental genetic markers

Genetic sex determination using the fraction of reads mapping to X and Y chromosomes showed that San Teodoro 3 was male while San Teodoro 5 was female, in agreement with the morphological results (Table S5, SI Appendix). Analysis of kinship indicated that the two individuals were not closely related (Table S6). The mitochondrial haplogroup of both individuals is U5b2b (Table S5), one of the most common mitochondrial haplogroups found in post-LGM hunter-gatherers from Europe^13,14^, and likely associated with the resettlement of Europe after the LGM^3^. The San Teodoro individuals belong to the same subclade as the 14,180-13,780 year-old Palaeolithic (Late Epigravettian) individual Villabruna from Northern Italy, which represents the most diverged haplotype (Figure S8, Table S7). Analysis of divergence times estimated the age of haplogroup U5b2b to be around 23 kya (from 19,137 to 27,984), in accordance with previous results^14^. Furthermore, the San Teodoro lineage is closely related with the Paglicci 71 lineage (dating back to around 18 Kya^13^) from Apulia (Italy), which refers to a previous phase of the Epigravettian (Evolved Epigravettian).

The Y-chromosome haplogroup determined for San Teodoro 3 is I2a2 (Tables S5, 8 and Figure S9), a subclade of haplogroup I2 which is common among European hunter-gatherers, and possibly originated in southern Europe during the LGM before becoming widespread in Europe during the Neolithic^15,16^. This haplogroup shows high frequency in present-day populations in the Balkan peninsula, in particular within the Serbian population^17^.

#### Genetic structure of European Hunter-Gatherers

For the purpose of analyzing the genetic diversity of European hunter-gatherers after the LGM, we merged the data from San Teodoro with a panel of previously published hunter-gatherer individuals (Palaeolithic and Mesolithic) from across Europe (Figure 1a). To visualize the genetic structure of the ancient individuals, we performed multidimensional scaling on a distance matrix derived from pairwise identity-by-state (IBS) sharing of alleles among individuals (Figure 1b, Figure S10). The two San Teodoro samples fell broadly within the genetic diversity of individuals from western and southern Europe, previously termed “Western hunter-gatherers” (WHG)^18^ (Figure 1b, Figure S10, S11). Within this group of individuals, San Teodoro clustered most closely with another Late Epigravettian individual from Sicily, Oriente C, and with the Late Upper Palaeolithic/Mesolithic individuals from Grotta Continenza (central Italy), indicating a common gene pool for post-LGM hunter-gatherers from the Italian peninsula from ~14,000 years ago onwards.

**Figure 1.**
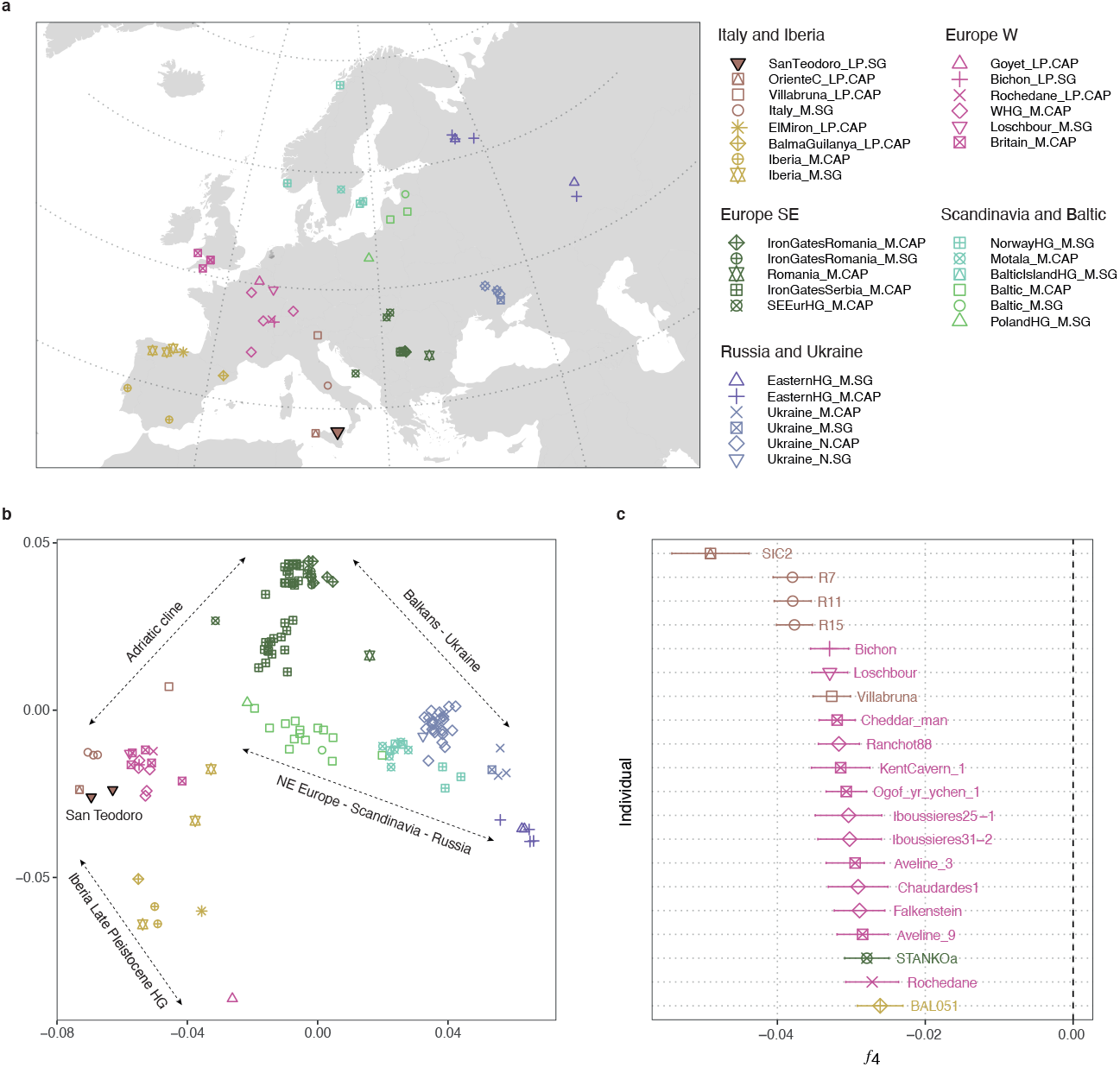
Genetic clustering of European post-LGM hunter-gatherers. **a,** Geographic location of new individuals from San Teodoro and previously published post-LGM hunter-gatherer individuals; **b,** Multidimensional scaling (MDS) of 138 hunter-gatherer individuals (>50,000 SNPs covered) based on a pairwise identity-by-state (IBS) allele sharing. Major ancestry clines are indicated with dotted arrows. **c,** Genetic affinities with the San Teodoro individuals. Shown are point estimates and standard errors for statistic *f*_4_(Mbuti, Test; SanTeodoro_LP.SG, DevilsCave_N.SG), which measures allele sharing of a test individual with San Teodoro compared to Neolithic hunter-gatherers from East Asia. More negative statistics indicate higher genetic affinity with San Teodoro. The top 20 individuals with the highest affinity are shown.

The clustering results are also confirmed by *f*_4_-statistics of the form *f*_4_(Mbuti, Test; SanTeodoro_LP.SG, DevilsCave_N.SG), which measures the excess of genetic drift a test individual shares with San Teodoro (from here on merged into a single population due to their high genetic similarity) compared to an outgroup (East Asian hunter-gatherers from Devil’s Gate cave). Results showed that WHG individuals, in particular those from the Italian peninsula, shared the highest amounts of genetic drift with San Teodoro. The highest affinity was observed for Oriente C (SIC2), an individual related to the same cultural sphere of San Teodoro (Late Epigravettian), with a similar age from the same geographic region (Sicily)^19^ (Figure 1c, Table S9).

Using *f*_4_-statistics of the form *f*_4_(Mbuti, Test; HG Italy, San_Teodoro_LP.SG), we next tested whether San Teodoro forms a clade with Italian hunter-gatherer groups, consisting of both Late Upper Palaeolithic (Oriente C, and Villabruna) and Late Upper Palaeolithic/Mesolithic (Continenza), to the exclusion of other test groups. While a clade-like relationship cannot be rejected for the Sicilian hunter-gatherer from Oriente C, significant statistics were observed in tests with the other two groups (Figure S12, Table S10). For the central Italian individuals from Continenza, most post-LGM individuals showed significantly increased shared genetic drift compared to San Teodoro. The consistent magnitude of the statistics across individuals with a wide geographic distribution, and chrono-cultural differentiation, suggests possible gene flow between the ancestors of most post-LGM Europeans and Continenza, after their divergence from San Teodoro. Interestingly, the only individual with evidence for higher affinity with San Teodoro than with Continenza was the ~33,000 year-old pre-LGM individual Paglicci133 from Apulia in southern Italy (*f*_4_(Mbuti, Paglicci133; Italy_M.SG, San_Teodoro_LP.SG) = 0.003, Z=2.6; Figure S12, Table S10). Paglicci 133 refers to Gravettian which is considered the cultural root (the Gravettian) from which the San Teodoro one (the Epigravettian) was derived in the following several millennia. This suggests possible gene flow involving ancestry related to pre-LGM hunter-gatherers in southern Italy, as previously observed on the Iberian Peninsula^18^, as a possible alternative explanation. A clade-like relationship was also rejected for the relationship of San Teodoro and the ~14,000 year-old northern Italian individual from Villabruna. In contrast to the results obtained with Continenza, in this analysis the individuals from western Europe mostly shared more genetic drift with San Teodoro, whereas individuals from eastern Europe shared more genetic drift with Villabruna (Figure S12, Table S10). These results were also reflected in the genetic clustering, where Villabruna was shifted away from the Italian and west European individuals on a cline towards individuals from the Balkans (Iron Gates, Figure 1b), suggesting further sub-structure among individuals of the “Villabruna cluster”.

The genetic diversity of European post-LGM hunter-gatherers has previously been described in terms of a west-to-east cline anchored by two major ancestry groups of “western hunter-gatherers” (WHG) and “eastern hunter-gatherers” (EHG)^15^, with some contributions from late Pleistocene hunter-gatherer ancestry in the Iberian Peninsula^20^. In the MDS in Figure 1b, individuals were maximally differentiated in four distinct ancestry clusters, namely: (i) hunter-gatherers from the Italian peninsula (including San Teodoro) on the left, (ii) late Pleistocene hunter-gatherer ancestry (represented by Goyet Q-2) at the bottom, (iii) individuals from the Balkans (Iron Gates) at the top, and (vi) EHG from Russia on the right. The remaining individuals were aligned across a number of distinct genetic clines, broadly related to their geographical location, including (Figure 1b): (i) an “Adriatic” cline, linking the Italian peninsula to the Balkans; (ii) an “Iberia late Pleistocene HG” cline of Goyet Q-2 related ancestry in individuals from the Iberian peninsula,; (iii) a “Balkans-Ukraine” cline; and (vi) a “NE Europe - Scandinavia - Russia” cline, linking individuals from the Baltic to Scandinavia and EHG. Furthermore, we found evidence for temporal genetic structure within geographic regions. This was most notably at Iron Gates, where the earlier individuals from Serbia (before 9000 BP), together with the neighbouring individuals from Romania, form one of the four most differentiated clusters (at the top in Figure 1b), whereas later individuals from Serbia (after 9000 BP) were shifted towards those from north-eastern Europe and the Baltics (Figure 1b, Figure S13).

Motivated by these observations, we used *qpAdm* to infer ancestry proportions of European hunter-gatherers in a four-way model using representatives of the maximally differentiated genetic clusters as sources. The results recapitulated many of the features observed in the genetic clustering. In individuals from western Europe, ancestry related to Italian hunter-gatherers predominated, with varying contributions of late Pleistocene hunter-gatherer (Goyet Q-2) ancestry (Figure 2, Figure S14, Tables S11-S14). Late Pleistocene ancestry peaked in the Iberian Peninsula, with inferred ancestry proportions as high as 71% (El Miron), consistent with previous results^20^. An apparent influx of Balkan hunter-gatherer related ancestry was observed in the two most recent individuals from northern Iberia (La Brana, Los Canes), suggesting gene flow between distinct ancestry groups towards the end of the hunter-gatherer dominion in western Europe^21^. In eastern Europe, ancestry related to Balkan hunter-gatherers was most abundant, with additional contributions of Italian or EHG ancestry, in line with the geographic origin of the individuals. Genetic continuity could be rejected for many of the later Balkan hunter-gatherers from Iron Gates, which showed evidence for admixture with groups harboring both Italian- and Goyet Q-2-related ancestry (Figure S14; Tables S11-S14). Finally, EHG-related ancestry was found at highest proportions in individuals from Ukraine, as well as Scandinavian hunter-gatherers^22^. Once again, we observed temporal stratification in ancestry suggestive of local genetic transformations, with an increase in Balkan hunter-gatherer related ancestry in Ukrainian individuals after 10,000 BP^15^ (Figure S14). Taken together, these results document previously underappreciated complexity in the fine-scale genetic structure of post-LGM hunter-gatherers in Europe.

**Figure 2.**
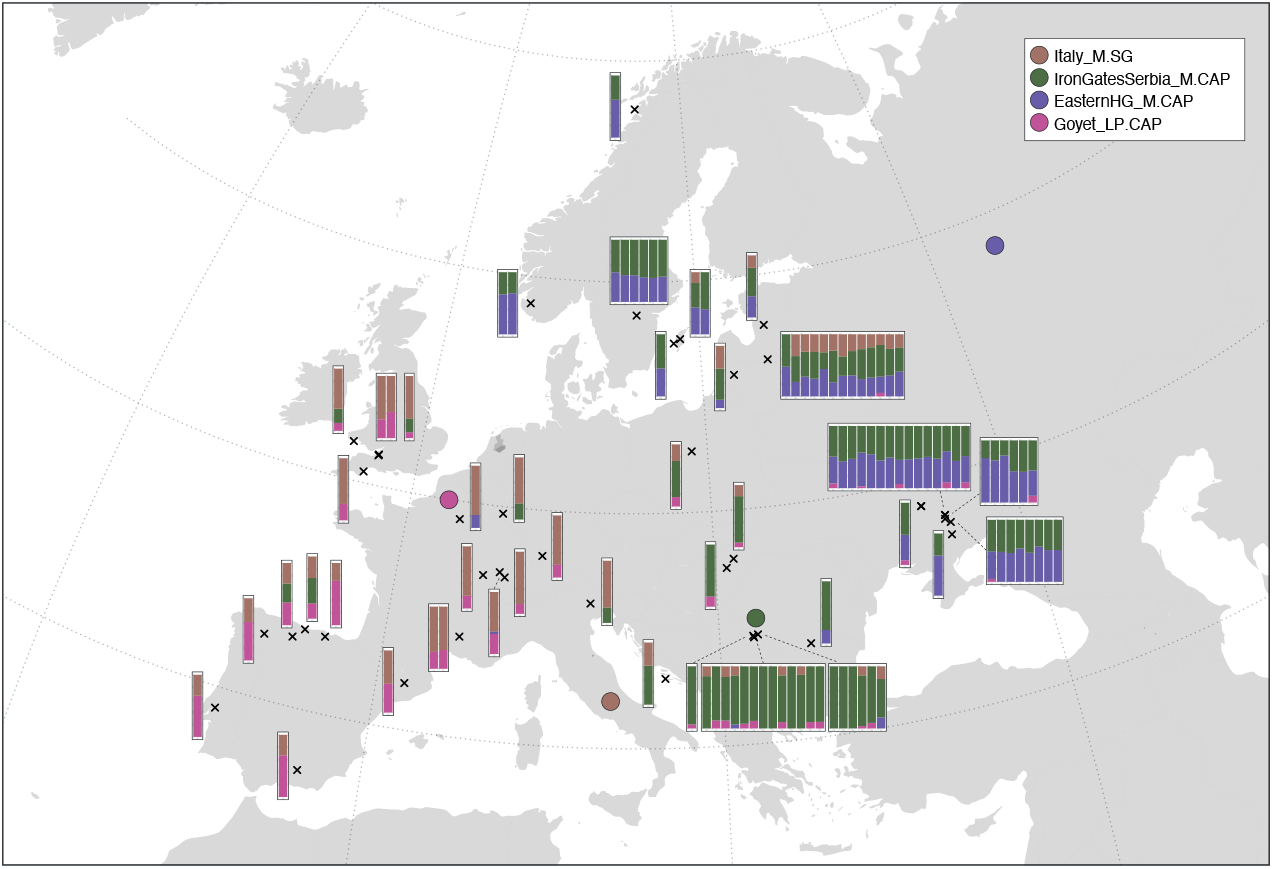
Spatial distribution of hunter-gatherer ancestry clusters. Map showing geographic locations (black crosses) and ancestry proportions (bar plots) of post-LGM hunter-gatherers, inferred using *qpAdm*. Individuals were modelled using four source groups, representing major post-LGM lineages identified in this and previous studies. Approximate geographic locations of source groups are indicated with colored symbols.

### Dental calculus

#### Diversity of the oral microbiome

We sequenced ancient DNA extracted from dental calculus in order to characterize the oral microbial communities of the two Upper Palaeolithic hunter-gatherers. We obtained totals of 28,732,940 (San Teodoro 3) and 32,249,586 (San Teodoro 5) sequencing reads, which were subjected to metagenomic classification using Kraken^23^ and KrakenUniq^24^, using a database of all reference sequences from the NCBI RefSeq database (February 2017), followed by species-level abundance estimation using Bracken^25^. This resulted in a final dataset containing 1,639,575 (5.71%) classified reads in San Teodoro 3, and 2,431,172 (7.54%) classified reads in San Teodoro 5. We first characterized the broad microbial composition of the two San Teodoro individuals (Tables S15-S16). Both were abundant in *Actinomyces, Streptococcus* and *Propionibacterium*, genera typically associated with the oral microbiome (Figure 3, Figures S16-S18, Table S15). San Teodoro 3 was also abundant in *Olsenella*, known to cause endodontic infections, while San Teodoro 5 was abundant in *Aggregatibacter* and *Neisseria*, both a part of the normal oral microbiome. *Lautropia* and *Ottowia* were found to be more abundant in samples from Neolithic farming and later contexts. *Mesorhizobium* is a soil bacterium and probably an environmental contaminant, though also present at low abundance in some modern oral samples.

**Figure 3.**
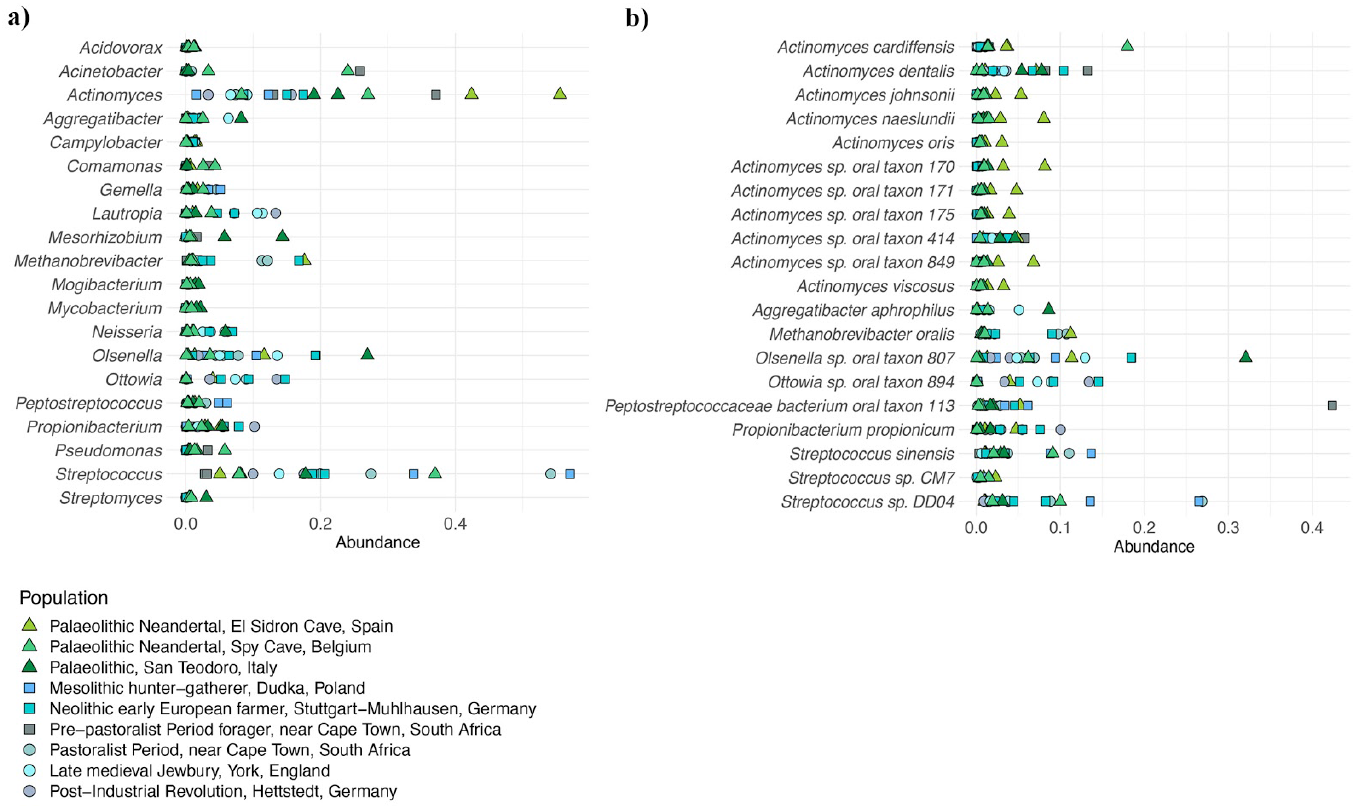
Metagenomic composition of ancient dental calculus. **a)** Abundance of the top 20 most abundant genera in ancient samples (Table S15). **b)** Abundance of the 20 most abundant species in ancient samples (Table S16).

Sequences of putative microbial origin associated with the oral microbiome showed characteristic ancient DNA damage patterns, supporting the authenticity of the data (Figure S15).

We analysed the ancient microbiome data from San Teodoro in the context of a reference dataset that comprised a total of 1,400 modern samples covering eight oral sites and four other human microbiomes, as well as 17 previously published ancient calculus samples^26^ (Table S17). To minimize possible impacts of microbial species from environmental contamination in the ancient samples, we restricted all compositional analyses to sets of microbial species with a minimum of 1,000 unique kmers classified in five or more modern samples, using KrakenUniq^24^. For analyses involving all modern and ancient samples (species set “all”), microbial species identified in all modern metagenomes were included (Table S18). For analyses restricted to the oral microbiomes (species set “oral”), only microbial species identified in modern samples from oral sites were included (Table S18).

We investigated differences in microbial composition using both unsupervised (Principal Component Analysis (PCA), dendrograms) and supervised (Discriminant analysis of principal components (DAPC)) clustering methods. All methods consistently showed samples clustering into communities according to their site of isolation (Figure 4a and Figures S19-S28). Samples from the retroauricular crease and the external naris clustered together, with some overlap of vaginal samples, consistent with previous results^27^. The ancient calculus samples clustered nearest to the plaque or the gingiva samples and were broadly divided according to mode of subsistence: Palaeolithic, foragers, and farmers (Figure 4a and Figures S19-S28). In the PCA, the more recent samples from the introduction of farming and forward were placed nearer to the modern samples than the Palaeolithic and pre-pastoralists (Figures S19-S22) and one of the post-Industrialization samples (IR_13234) clustered with modern samples. Allowing K-means to decide the groupings of all samples before DAPC, all ancient samples were found to cluster together. An exception was one of the Neandertal individuals from Spy cave (SPY1), which clustered with samples from retroauricular crease and the external naris and has previously been found to be contaminated with modern environmental DNA^27^. Modern samples are largely clustered according to their isolation source, though split into smaller groups (Table S19). When analysing only the oral samples and corresponding species list, we observed further finer-scale structure. Groups identified by K-means separated all Palaeolithic samples and one pre-pastoralist forager from later hunter-gatherer, Neolithic, and post-Neolithic samples (Figures S25-S26 and Table S20). With the exception of one ancient sample from the Post-Industrial Revolution period (IR_13234) which clustered with modern oral samples, both clusters of ancient samples were clearly distinct from the modern microbiomes. Restricting to species identified in modern oral samples also diminished the influence of contamination in SPY1, which clustered together with the other Palaeolithic individuals in this analysis.

**Figure 4.**
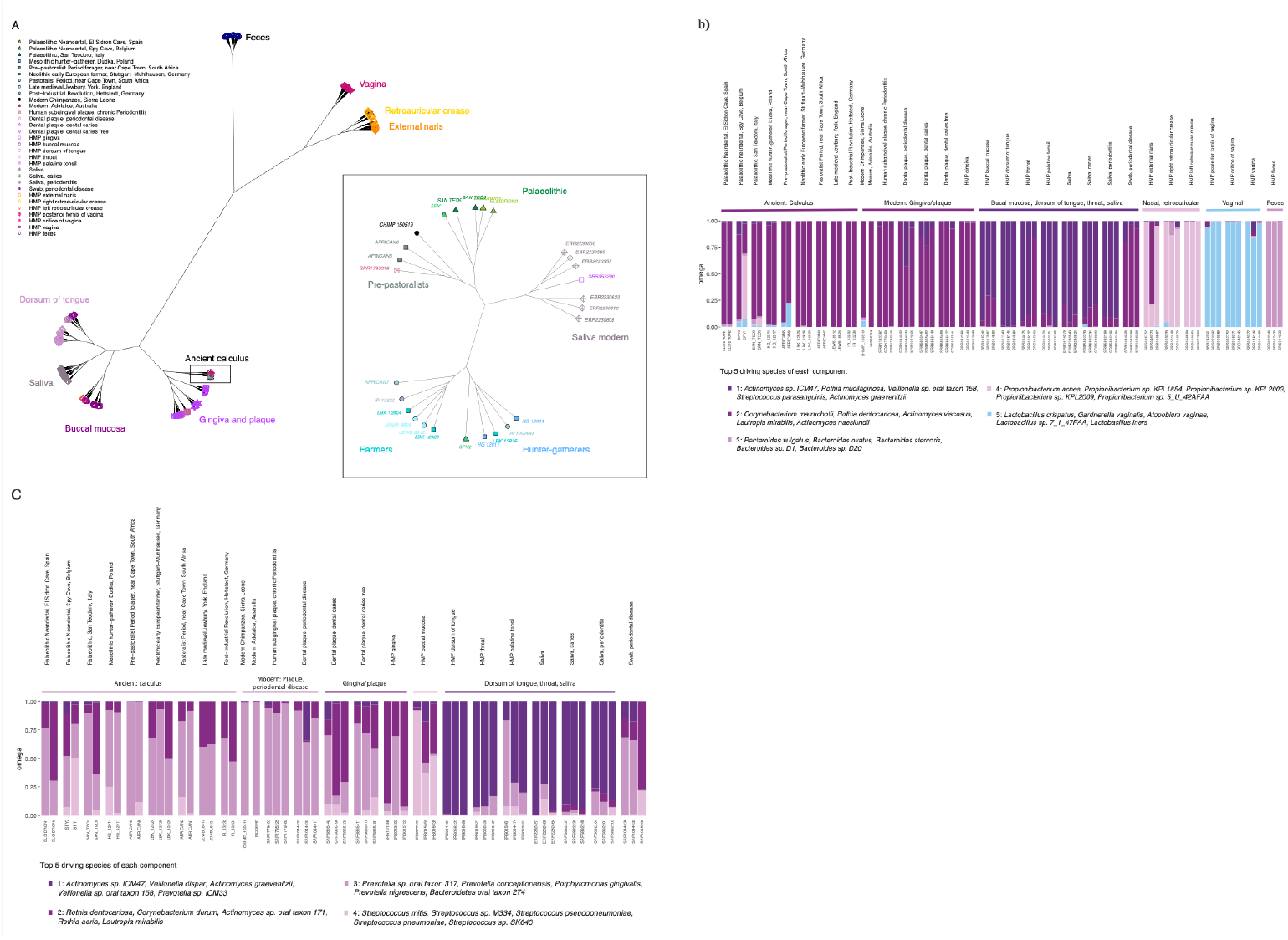
Clustering of ancient microbiomes. **a)** Dendrogram created using the species set ‘all’. Green inset shows structure among ancient samples only. **b)** Bar plots showing Grade of Membership (GoM) created using species set ‘all’, for k=5 components. **c)** GoM including only oral samples and created using the species set ‘oral’, k=4 components.

To further investigate the microbial species distinguishing the different microbiomes and between ancient and modern samples, we performed a Grade of Membership (GoM) analysis^28^. The analysis was run with two to nine components (k=2 to k=9), with both all human microbiome samples and only oral samples. A random subsampling of three samples per site is shown for k=5 for the full microbiome analysis and k=4 for the oral microbiome (Figure 4b, c, Figures S29-S30). In the full microbiome analysis using two components, samples from oral sites and feces were separated from those of the external naris, retroauricular crease, and vaginal sites. Among the ancient samples, SPY1 showed a substantial fraction of its GoM from a component maximized in samples from the external naris, retroauricular crease, consistent with the aforementioned contamination with non-oral microbial communities^26^. At k=3, vaginal sites were separated from retroauricular creases and external naris, which are characterized with species of the genus *Propionibacterium* commonly associated with skin microbiome^29^. A distinction between oral sites was first observed at k=5, with the ancient calculus samples more similar to gingiva and plaque, consistent with their sampling location (Figure 4b). Higher numbers of components revealed additional substructure within microenvironments of the same body site, such as the buccal mucosa at k=7 which was associated with species of the genus *Streptococcus* such as *Streptococcus mitis* and *Streptococcus pneumoniae*. At k=9, we observed a division between healthy and periodontal gingiva or plaque samples, associated with *Prevotella* sp. and *Porphyromonas gingivalis* as the driving species for periodontal samples (Figure 4b, Figure S29). Restricting to species found in the oral microbiome only, we found that three components (k=3) separated the different oral compartments of gingiva/plaque, buccal mucosa, and dorsum of tongue and saliva. The ancient calculus samples were similar to modern gingival and plaque samples, and their component was associated with species such as *Corynebacterium matruchotii, Propionibacterium propionicum*, and *Actinomyces naeslundii*. At k=4, healthy and periodontal gingiva/plaque samples were distinguished. Clusters with high proportions in healthy samples were associated with species such as *Rothia dentocariosa, Corynebacterium durum, Rothia aeria*, and *Lautropia mirabilis*. Samples with periodontal disease on the other hand were characterized by a component including gram-negative, anaerobic bacteria known to show increased abundance with periodontal disease^30^, such as *Prevotella conceptionensis, Porphyromonas gingivalis* and *Prevotella nigrescens*. Interestingly, most of the ancient calculus samples were also highly abundant in the periodontal component (Figure 4c, Figure S30).

We further characterized differential microbial abundance among the samples using ALDEx2, a method developed for the analysis of compositional data^31,32^. Applying the method to our dataset, we found species including *Methanobrevibacter oralis, M. smithii, Olsenella sp. Oral taxon 807* and *Peptostreptococcaceae bacterium oral taxon 113* among those with significantly higher abundance in the ancient samples (Table S21-S24, Figure S31-S32). On the other hand, species of the genera *Veillonella, Streptococcus, Haemophilus, Rothia, Porphyromonas, Campylobacter* and *Neisseria* were significantly more abundant in the modern samples. We found that gingiva/plaque sites were overall depleted in *Streptococcus* species, particularly in samples from individuals with periodontal disease*. Capnocytophaga* species were more abundant in gingiva/plaque samples, whereas *Porphyromonas gingivalis* and *Treponema denticola* were more abundant in samples with periodontal disease and ancient calculus. Interestingly, *T. vincentii* and *Prevotella* species were also more abundant in modern periodontal samples, but not in ancient samples. We did not find any significant differentiation between pre- and post-Neolithic ancient samples (Table S25).

#### Identification of dietary and human proteins

We used palaeoproteomics to identify dietary and human proteins present in the dental calculus of the San Teodoro individuals (Table 1, Table S26 and S27). To support the endogenous origin of the identified proteins, we calculated the rate of deamidation for asparagine and glutamine, a spontaneous form of hydrolytic damage consistently observed in ancient samples^33^. Both the samples show an advanced rate of deamidation, compatible with the authenticity of the generated data (Figure 5).

**Table 1:** Dietary proteins detected in the San Teodoro 3 and 5 dental calculus samples. * The identification of wild extinct species: *Bos primigenius* and *Equus hydruntinus*, is supported by the observation of tandem MS spectra confidently matching peptides specific to the proteome of domestic species of the same genus: *Bos taurus* and *Equus caballus/Equus asinus*, respectively. Bone remains from *Bos primigenius* and *Equus hydruntinus*, dated to the Late Upper Palaeolithic at ~15ky cal. BP, have been found in the San Teodoro cave and in other Late Epigravettian sites in Sicily^36^.

|  | Accession numbers | Protein name | Species | Interpretation | Total peptides | Specific peptides | Unique + razor sequence coverage [%] | Unique sequence coverage [%] | Sequence length | Number of MS/MS |
| --- | --- | --- | --- | --- | --- | --- | --- | --- | --- | --- |
| **San Teodoro 3** |  |  |  |  |  |  |  |  |  |  |
| Animal | P02453 | Collagen Type I Alpha 1 Chain | *Bos taurus* | *Bos primigenius* * | 30 | 1 | 24,1 | 1,23 | 1463 | 84 |
|  | P02465 | Collagen Type I Alpha 2 Chain | *Bos taurus* | *Bos primigenius* * | 21 | 4 | 15,2 | 5,64 | 1364 | 56 |
|  | P04258/XP_005675926.1/W5Q4S0 | Collagen Type III Alpha 1 Chain | *Bos taurus/Capra hircus/Ovis aries* | *Bos primigenius* * | 4 | 2 | 4,3 | 2,86 | 1467 | 6 |
| Fish | KTF73577.1 ; KTG05890.1 | hypothetical protein cypCar_00002572; hypothetical protein cypCar_00045321 | *Cyprinus carpio* | *Cyprinus carpio* | 5 | 1 | 3 | 0,74 | 1352 | 8 |
| **San Teodoro 5** |  |  |  |  |  |  |  |  |  |  |
| Animal | P02453 | Collagen Type I Alpha 1 Chain | *Bos taurus* | *Bos primigenius* * | 38 | 1 | 29,9 | 1,23 | 1463 | 127 |
|  | P02465 | Collagen Type I Alpha 2 Chain | *Bos taurus* | *Bos primigenius* * | 23 | 5 | 12,5 | 6,82 | 1364 | 63 |
|  | F1SFA7/A0A1S7J1Y9 | Collagen Type I Alpha 2 Chain | *Sus scrofa/Sus scrofa domesticus* | *Sus scrofa* | 5 | 1 | 4,5 | 1,1 | 1135 | 54 |
|  | A0A287BLD2/A0A1S7J210 | Collagen Type I Alpha 1 Chain | *Sus scrofa/Sus scrofa domesticus* | *Sus scrofa* | 4 | 1 | 4 | 1,86 | 1451 | 85 |
|  | F6RTI8/B9<br>VR89 | Collagen Type I Alpha 2<br>Chain | *Equus*<br>*caballus/Equus*<br>*asinus* | *Equus*<br>*hydruntinus\** | 2 | 2 | 2 | 1,99 | 1357 | 41 |
|  | XP_0179203<br>82.1/W5P48<br>1/XP_01198<br>3013.1 | Collagen Type I Alpha 1<br>Chain | *Capra*<br>*hircus/Ovis*<br>*aries/Ovis aries*<br>*musimon* | *Capra ibex* | 5 | 2 | 5,1 | 2,44 | 1474 | 127 |
|  | F1N2Y4/W5<br>Q4S0/XP_0<br>05675926.1 | Collagen Type III Alpha<br>1 Chain | *Bos taurus/Ovis*<br>*aries/Capra*<br>*hircus* | *Bos*<br>*primigenius\** | 6 | 3 | 10 | 4,97 | 1047 | 11 |
| Legumes | 5GYL | Legumin-like | *Cicer arietinum* | *Lathyrus* | 15 | 6 | 39,5 | 17,95 | 479 | 27 |
|  | XP_0044930<br>35.1 | Vicilin-like | *Cicer arietinum* | *Lathyrus* | 10 | 6 | 22,2 | 15,6 | 455 | 12 |
|  | XP_0044937<br>80.1 | Legumin A-like isoform<br>X1 | *Cicer arietinum* | *Lathyrus* | 7 | 4 | 18,7 | 13,51 | 518 | 18 |
|  | XP_0044951<br>00.1 | Legumin J-like | *Cicer arietinum* | *Lathyrus* | 7 | 5 | 17,3 | 12,6 | 532 | 10 |
|  | XP_0044967<br>03.1 | Provicilin-like | *Cicer arietinum* | *Lathyrus* | 4 | 3 | 7 | 5,73 | 558 | 4 |
|  | XP_0044967<br>04.1 | Vicilin-like | *Cicer arietinum* | *Lathyrus* | 2 | 2 | 7,7 | 6,4 | 377 | 5 |
| Fish | KTF73577.1<br>;<br>KTG05890.<br>1 | hypothetical protein<br>cypCar_00002572;<br>hypothetical protein<br>cypCar_00045321 | *Cyprinus carpio* | *Cyprinus*<br>*carpio* | 5 | 1 | 4 | 0,9 | 1352 | 9 |
|  | ABY71227 | Collagen Type I Alpha 1<br>Chain | *Scyliorhinus*<br>*canicula* | *Scyliorhinus*<br>*canicula* | 2 | 2 | 2,1 | 2,1 | 1237 | 3 |

**Figure 5:**
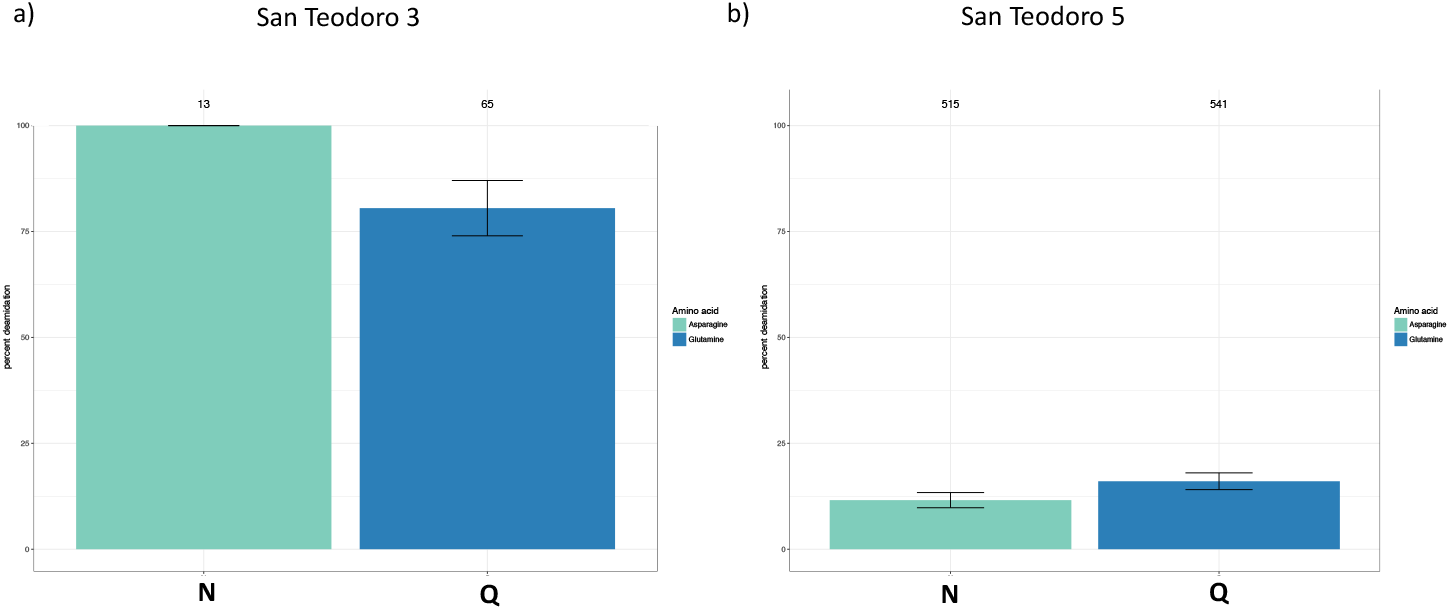
Proteins deamidation. Overall percentage of deamidation for asparagine (N) and glutamine (Q) amino acids for the proteins found in the dental calculus samples: a) San Teodoro 3; b) San Teodoro 5. Numbers above each bar represent the number of peptides used for the analysis and the error bars represent standard deviation

To avoid spectra misinterpretation, we included all common modifications that could affect ancient proteins, such as deamidation, as variable modifications in the analyses (see materials and methods) and all the spectra associated has been manually inspected, validated, and annotated. Compared to San Teodoro 3, San Teodoro 5 shows a lower rate of damage and a higher number of confidently identified peptides and proteins, suggesting a different state of protein preservation for the two San Teodoro dental calculus samples (Figure 5, Table 1, Table S26, S27). In San Teodoro 3, the human proteins identified are limited mostly to collagen^34^. In San Teodoro 5, a total of 22 human proteins are observed (Table S26), most of which were associated with immune response, as previously observed^10^.

The food proteins provide an unprecedentedly detailed insight into the diet of the San Teodoro individuals. While stable isotope analyses previously could only suggest a relevant consumption of animal proteins, typical of hunter-gatherers^35^, palaeoproteomic analysis identified some of the plant and animal taxa each individual consumed (Table 1). Specifically, in both calculus samples from San Teodoro 3 and 5 (Table 1 and Table S27) we identified bovine Collagen Type I Alpha 1 and 2, arguably from aurochs (*Bos primigenius*), the only bovine species known to live in Sicily during the Late Upper Palaeolithic. This observation was also supported by the recovery of DNA sequences originating from the genus *Bos* in the metagenomic data of San Teodoro 3 (Table S28). For similar reasons, collagen proteins found in San Teodoro 5 and confidently assigned to pig (*Sus scrofa/Sus scrofa domesticus*) and equine (*Equus caballus/Equus asinus*) should be attributed to the wild species *Sus scrofa* and *Equus hydruntinus* (Table 1 and Table S27). Our results therefore conclusively support the consumption of aurochs, wild boar and European wild ass, as previously suggested by the recovery of bone remains from these taxa in the faunal record of many Late Epigravettian Sicilian deposits^36^ including San Teodoro cave (SI Appendix).

The palaeoproteomic results also showed the consumption of fish in both samples (Table 1 and Table S27). In San Teodoro 3, only one freshwater fish taxon, common carp (*Cyprinus carpio*), was identified, while in San Teodoro 5, both freshwater and marine fishes were found: *Cyprinus carpio* and the small, shallow-water shark *Scyliorhinus canicula*. We caution the peptides recovered could also match other close taxa, whose proteome sequences are not yet publicly available. While cyprinids are frequently recorded in Western European Upper Palaeolithic deposits, including Epigravettian sites in continental Italy^37^, no evidence of this group has been previously reported from Palaeolithic deposits in Sicily. The presence of freshwater and marine fish in the diet of the two individuals from the San Teodoro cave is fully coherent with its geographic position (a few hundred meters away from the streams Inganno and Furiano, and from the marine coast, Figure S2) and in accordance with the archaeological record know in Sicily and the Mediterranean area for the Upper Palaeolithic^38–40^.

In San Teodoro 5 six seed storage proteins matches modern chickpea (*Cicer arietinum*) (Table 1 and Table S27). Chickpea is one of the crops present in primeval agriculture in the Near East and Europe. The first archaeological findings of this species are associated with the Pre-pottery Neolithic site (El-Hemmeh) in Jordan dated back 11,100-10,610 BP^41^. From the Near East, the chickpea spread to south-eastern Europe with the Neolithic transition^41^. There is no evidence of chickpea consumption during the Upper Palaeolithic in southern Europe. At the end of the Iron Age in the Mediterranean regions, the main spontaneous pulses were the red pea (*Lathyrus cicera*) and the grass pea (*Lathyrus sativus*). Their consumption was attested in Spain during the Upper Palaeolithic^42^, however so far there is no palaeobotanical evidence of the presence of this group of wild plants in the Sicilian Upper Palaeolithic. At the time our analysis was performed, the plant proteins we identified were not among those whose sequence was known for the genus *Lathyrus*. Based on the archaeological context the samples originate from, we exclude the consumption of chickpeas and consider instead the exploitation of wild pulses highly plausible, possibly of the *Lathyrus* genus. All the plant proteins we identified are seed storage proteins, and their consumption may have occurred after the transformation of the seeds into flour. The use of flours from wild plants is documented in other contexts of the Upper Palaeolithic^43^.

Finally, in San Teodoro 5 we identified Collagen Type I Alpha 1 matching domesticated ovicaprid species: *Capra hircus/Ovis aries/Ovis aries musiman*, none of which is likely to be present in the archaeological context of San Teodoro and in the other Epigravettian sites in Sicily. Mouflon (*Ovis gmelini*), the wild species closest to sheep, is not present in Late Upper Palaeolithic in Southern Europe, while ibex (*Capra ibex*), the wild species closest to goat, is well documented during the Late Glacial in Southern peninsular Italy (the southernmost and closest documentation to Sicily is in the Grotta del Romito in Northern Calabria)^44^ and never inhabited in Sicily^45^. Then we interpret the identification of ovicaprid collagen as a possible evidence of ibex exploitation.

## Discussion

In recent years, rapid progress has been made in the recovery of DNA from ancient human remains. These data have revolutionized our understanding of human demographic history, and the processes that shaped genetic diversity in the past. However, this focus on the analyses of ancient human DNA has, until recently largely neglected important other aspects of the human past, such as inferences about lifestyle and behaviour of prehistoric individuals. In this study, we combined ancient genomic, metagenomic and palaeoproteomic data to characterize ancestry, diet and microbial environment of an Upper Palaeolithic hunter-gatherer community from Sicily.

The genetic ancestry of the Palaeolithic population of San Teodoro fell broadly within the variation associated with hunter-gatherer individuals from Europe, in particular those of other central-southern Italian Late Upper Palaeolithic/Mesolithic individuals: Oriente C and Grotta Continenza. The high genetic affinity among the Sicilian HGs suggests that a founder effect may have played a key role in determining the genetic makeup of the post-LGM Sicilian population. We observed several genetic clusters and clines in European HGs, associated with the geographical localization of the individuals studied. Among them, our human genetic results highlight the presence of an Italian genetic cluster which likely played a key role during the post-LGM resettlement of western Europe. These results are in accordance with the contraction of animals and plants species in southern areas of Europe, during the LGM^1^ and the following expansion from these glacial refugia to northern and central areas of Europe^3,13^ from 18 kya BP, thanks to the rapid climatic amelioration. On a continental scale, our results suggest that geographic clines and isolation-by-distance played an important role in shaping European hunter-gatherer diversity. Nevertheless, we also find evidence for local transformations and possible migrations in regions including northern Iberia, the Balkans and Ukraine.

The integration between the metagenomic and the proteomic analysis of dental calculus provides a detailed insight into the diet and the composition of the oral microbiome of the human hunter-gatherers during the Upper Palaeolithic. Specifically, we find, in line with previous results^27^, that ancient calculus microbiomes broadly cluster according to the mode of subsistence, which is also true for these Epigravettian individuals. The oral microbiomes and the several oral animal proteins found in San Teodoro individuals are consistent with the well-founded meat-rich diet hypothesis already inferred by the rich archaeological (faunal) record from Late Upper Palaeolithic contexts and further supported by the high nitrogen stable isotope ratio values identified in several Late Upper Palaeolithic individuals from Sicily^35^. This result is also in accordance with the protein data where a lack of plants proteins was observed in the samples analysed. In fact, only the consumption of legume seed storage proteins was identified in San Teodoro 5 was. It should be noted that the absence of specific food proteins does not necessarily mean that a particular food resource was not regularly ingested. However, the differentiation between the San Teodoro and early farmer oral microbiomes and the lack of plant proteins found in the samples analysed could suggest low plant intake in this Upper Palaeolithic community. Still, this result confirms that the dietary habits of Epigravettian hunter-gatherers included some plant foods, as suggested in other archaeological contexts in Northern and Central Italy^39,46^.

The identification of proteins matching the genus of Capra, which could possibly belong to ibex, opens speculations on the dynamics of human populations of Sicily in the Late Glacial and can contribute to the study of mobility of Upper Palaeolithic hunter-gatherers between the island and the Italian peninsula or some trade between Calabria and Sicily. So far there is no archaeozoological and palaeontological evidence supporting the presence of ibex in Upper Palaeolithic Sicily^47^, instead, this species was very common on the mountains of nearby Calabria during the Upper Palaeolithic, for instance at the site of “Grotta del Romito”^44^. The hypothesized consumption of ibex by the San Teodoro 5 individual can either be an indicator of the movement of the San Teodoro 5 individual to (or from) peninsular Italy, even though an indirect supply cannot be ruled out as a consequence of transport of ibex meat from Calabria to Sicily by other Epigravettian hunter-gatherers. Either way, the results of the proteomic analysis provide important insights to advance hypotheses on the mobility of the Sicilian hunter-gatherers and on their contacts with the Italian peninsula. In addition, the individuals of San Teodoro are not the oldest evidence of human colonization of Sicily, which, on a radiometric basis and in accordance with current archaeological evidence, dates back to about a millennium earlier^48,49^. Therefore, they could be evidence of a further arrival of human groups in Sicily from Calabria where they might have eaten ibex meat.

On the other hand, the possibility of occasional or habitual movements towards Southern Calabria by the Epigravettian hunters of San Teodoro, with a short sea-crossing, cannot be ruled out. However, the hypothesis of continuous movements between Sicily and Calabria by Epigravettians could conflict with the undoubted diversity between the features of both the Calabrian stone assemblages (Grotta del Romito) and the lower Tyrrhenian side (Grotta della Serratura) compared to the Sicilian ones during the entire Late Epigravettian chrono-cultural range (about 13-10,000 uncal bp). It is well known that Sicily is one of the Epigravettian regions forming the articulated framework of Late glacial techno-complexes in Italy, which presents significant peculiarities in lithic production compared to the other southern Italian regions^50^. In this regard, the lack of archaeological evidence of an Epigravettian occupation in southern Calabria could explain the absence of contact between the Sicilian Late Paleolithic groups and those of the lower Tyrrhenian side, as the differences in lithic production seem to suggest.

In conclusion, on strictly archaeological grounds, the data relating to the ibex meat consumption by ST5 individual must leave open the problem about possible movements by the Late Paleolithic hunters between Sicily and the Italian peninsula, pending acquisition of new information.

The presence of marine and freshwater fish species fits with the framework of knowledge about the exploitation of the aquatic resources in the Mediterranean basin during the Upper Palaeolithic. Sea fish consumption evidence at San Teodoro is in accordance with what is already known for the Late Palaeolithic in Sicily^51^ and in other coeval contexts from the Mediterranean basin. In Sicily, an increase in marine exploitation has been observed in the Mesolithic due to a combination of sea level rising, population growth, and terrestrial resource depletion after the LGM^51^. However, only a few Upper Palaeolithic Sicilian sites provide sufficient stratigraphic and chronological information about the exploitation of aquatic resources^51^. Our results further confirm and illuminate the exploitation of marine and freshwater resources during Late Epigravettian, showing the important benefits of the proteomic approach to identify species often absent in the archaeological records of the ancient sites.

In conclusion, the -omics approach showed the maximum use of the environmental resources available to these hunter-gatherers, and also reveals a better picture of the ecology of Sicily at that time. Taken together with the aDNA results of the individuals, we demonstrate the value of investigating multiple classes of ancient biomolecules from the same individuals. We believe that the increase of genetic data in the near future will help to gain a more comprehensive picture of the Late Upper Palaeolithic and Mesolithic periods using both genetic information and cultural data to follow the dynamics of human population in Europe.

## Materials and Methods

### Archaeological sample material

Two individuals from San Teodoro cave in Sicily were sampled for aDNA analysis and lifestyle evaluation. For the human aDNA analysis the petrous bone was selected. For the lifestyle evaluation, dental calculus was analyzed by mass spectrometry-based proteomics and for the oral microbiome reconstruction by the metagenomic approach. Dental calculus was carefully removed from the tooth using a sterile periodontal scaler and around 20-30 mg were transferred into DNA LoBind Eppendorf tubes. Between one sampling and another, the periodontal tools were cleaned with bleach (concentration 5%) and after with ethanol (70%)^52^. Each sample was then divided into two different tubes, 22.8 mg (San Teodoro 3) and 9.9 mg (San Teodoro 5) for metagenomics analysis and about 7 mg each for proteomic analysis. All the molecular work was performed in DNA clean laboratory facilities at the Lundbeck Foundation GeoGenetics Centre, Globe Institute of the University of Copenhagen.

### DNA Extraction

The Allentoft^53^ protocol was used for the aDNA extraction from both ancient matrices. A starting amount of 150-400 mg of bone powder and 10-23 mg of dental calculus were added a pre-digestion buffer and incubated for 45 minutes at 37°C in order to remove the surface contaminants. Negative extraction controls were processed along with the samples. After centrifugation at 2000 g for 2 minutes, the supernatants were discarded and a new digestion buffer was added, then the samples were left for 24 hours at 37°C. The DNA extraction was performed on the digestion buffer by Silica powder. After centrifugation at 2000 g for 2 minutes, the supernatant was transferred into new tubes and 100 ml of silica suspension and 10 x volume of binding buffer was added. This solution was incubated at room temperature for 1 hour. Afterwards, the samples were centrifuged at 2000 g for 2 minutes and the supernatant was discarded. An additional 1 ml of binding buffer was added to the samples and the DNA was cleaned by ice-cold ethanol in a new tube. Finally, the DNA was eluted in 90 μl Qiagen EB buffer.

### NGS library preparation

DNA libraries for sequencing were prepared following a method proposed by Allentoft^53^. This method is divided into 4 steps: End-repair, Quick Ligation, Fill-in, and Indexing. In each step, a negative library control was included.

For the End-repair, around 20 μl of DNA extraction was used and NEB End-repair (module E6050L) Mix was added, according to the manufacturer’s instructions, to make a total of 25 μl. The solution was incubated at 12 °C for 20 minutes and 37 °C for 15 minutes. Before the ligation a purification step by Qiagen MinElute spin columns was performed and the DNA was eluted in 17 μl EB buffer.

In the Quick Ligation step, Illumina specific adapters^54^ were added to 15 μl of purified DNA by adding the NEB Quick Ligation Mix (module E6056L) and the solution was incubated at 20 °C for 15 minutes. Then, the mix was purified using Qiagen MinElute spin columns and the DNA was eluted in 20 μl Qiagen EB buffer.

For the Fill-in, 30 μl of NEB (module M0275L) reaction mix was added and incubated at 65 °C for 20 minutes and 80 °C for 20 minutes. The library was then quantified using IS8 and IS7 primers and SYBER green solution, according to manufacturer’s instructions. The quantification results were used to assess the optimal number of PCR cycles required for DNA library indexing. The indexing was performed by adding 1 μl of each primer (10 μM, inPE forward primer and indexed reverse primer) and 2X Kapa (following the manufacturer’s temperature instruction). After the indexing, the amplified DNA was purified by Qiagen MinElute Kit, eluted in 50 μl EB buffer, and quantified using an Agilent Bioanalyzer 2100.

The libraries were shotgun sequenced by the Illumina HiSeq 2500 and HiSeq 4000 platforms (81-bp single-read) at the Danish National High-Throughput DNA Sequencing Centre, University of Copenhagen, Denmark. For the run on Illumina HiSeq 4000, the library preparation followed the same steps previously described, but dual-indexed libraries construction and amplification were used.

### Bioinformatics pipeline

AdapterRemoval 1.5.2^55^ was used to remove the Illumina adapter sequences. Reads were mapped against the human reference genome build 37 with BWA 0.6.2^56^. Only reads with above 30 in mapping quality was kept using samtools 0.1.18^57^. Eventually, the PCR duplicate reads were removed by Picard MarkDuplicate http://broadinstitute.github.io/picard/. The mapping statistics for each sample analyzed are reported in Table S5.

The sex determination was performed by Skoglund and colleagues^58^ python script based on Rg parameter (Rg value lower than 0.016 is consistent with a female while result above 0.077 with a male). The results are reported in Table S5.

### DNA authentication

DNA contamination is one of the most serious problems in the aDNA study of samples from museum collections, for this reason, strict approaches were applied. First of all, the deamination pattern in the 5’ and 3’end (C to T transition and G to A, respectively) was assessed by mapDamage 2.0^59^ (Figure S2a) in the results obtained from both the biological matrices.

Moreover, two different methods were also used for the human aDNA authentication: the estimation of contamination in X-chromosome (for male sample) and the evaluation of the probability of mitochondrial DNA contamination (Table S5, Figure S7b). The X-chromosome contamination was evaluated just for the male individual (San Teodoro 3) applying ANGSD package^60^ following the commands suggested by the authors.

The mitochondrial contamination was evaluated for both samples by contamMix 1.0^14^ that it reports a Bayesian-based estimate of the posterior probability of the contamination proportion.

### Mitochondrial and Y-chromosome haplogroups assignment

For both samples, the mitochondrial haplogroup was assessed by a command line of haplogrep2^61^ (Table S5).

While for San Teodoro 3 (male sample) also Y-chromosome haplogroup was identified. Bcftools mpileup and bcftools call were used for the genotypes call with only bi-allelic SNP sites of the Y chromosome from the International Society Of Genetic Genealogy (ISOGG, http://www.isogg.org, version 10.107). By epa-ng we built a phylogenetic tree of Y chromosome sequences from the 1000 Genomes project, with maximum likelihood placement of ST3 (Figure S9).

### BEAST analysis

BEASTv1.8.4^62^ was used to reconstruct the phylogenies of the U5b mitochondrial haplogroup. BEAUti v1.8.4 was applied to generate the input file for BEAST analysis. We used the radiocarbon dates of the ancient samples (Table S7) as calibration points in the tree inference, and after analysis with jModelTest 2.1.10^63^ we applied the HKY substitution model^64^ with gamma plus invariant sites and strict clock with a prior of 2.2 10^-8^ μ/site/year^65^. The phylogenetic reconstruction was carried out on 32 ancient samples with MareuilLesMeaux1, haplogroup U5a2^13^, as outgroup. We used the Extended Bayesian Skyline Plot method for mitochondrial and the MCMC chains were run for 10^8^ states and sampled every 10^4^ states. MCMC runs were evaluated using Tracer (v1.6) (http://tree.bio.ed.ac.uk/software/tracer/). The resulting trees were annotated by TreeAnnotator v1.8.4 and visualized by FigTreev1.4.3 (http://tree.bio.ed.ac.uk/software/figtree/).

### Kinship analysis

We also determined the kinship relationship between the San Teodoro individuals inferring rates of identity-by-descent (IBD) sharing between pairs of individuals by the kinship coefficient estimator implemented in KING^66^. This is based on pairwise identity-by-state (IBS) sharing, obtained estimating the 2D-SFS for each pair using realSFS tool from the ANGSD package^60^. Expected values for the different estimators and selected degrees of relatedness are shown in Table S6.

### Ancestry analysis

Analyses of genetic ancestry of the San Teodoro individuals were carried out using a dataset including previously published Eurasian hunter-gatherer individuals (Table S29). Genetic similarity measures were estimated based on pairwise sharing of alleles identical-by-state (IBS). Population structure was visualized using multidimensional scaling on a distance matrix calculated as 1-p(IBS). Patterns of admixture and shared genetic drift were evaluated using *f*-statistics, calculated using Admixtools 5.0^67^. Ancestry proportions of post-LGM hunter-gatherers were inferred using *qpAdm*, using a set of 12 outgroups (Mbuti.SG, Yana_UP.SG, UstIshim, Sunghir_UP.SG, IronGatesRomania_M.SG, CHG_M.SG, Botai_EBA.SG, FunadomariJomon_N.SG, Kolyma_River, Zagros_EN.SG, SanTeodoro_LP.SG, EasternHG_M.SG).

### Dental calculus metagenomics analysis

The metagenomic part includes two ancient Upper Palaeolithic Italian calculus samples from San Teodoro. We also included the published calculus samples from Weyrich and colleagues^26^ and for modern comparison, all whole-genome sequenced Human Microbiome Project (HMP)^27^ samples, and six other oral microbiome studies, PRJNA383868, PRJNA230363, PRJNA255922, PRJNA396840, PRJEB14383, PRJEB24090, with a known source material. An overview of the included ancient samples is shown in Table S15.

All samples were trimmed with AdapterRemoval^55^ with a minimum length of 30 bp and quality above 20 on the phred scale. All ancient samples were run through an in-house pipeline conducting FastQC 0.11.5^68^, Kraken 1.0^23^, Bowtie2 2.3.2^69^ and mapDamage^59^. Kraken was also applied on all modern samples. Bracken^25^ was applied afterwards for species-level abundance estimation. Furthermore, KrakenUniq^24^ was used to filter for false positives and environmental contaminants of the ancient samples. The data was filtered by a species list including bacteria, fungi and archaea. Only species with above ten reads classified and at least 1000 unique k-mers, or twice as many unique k-mers as reads were included, if present in at least five modern samples. All other analysis was done in R version 3.5.1. For the distance-based methods, the transformation method for compositional data proposed by Gloor et al.^70^ was used, i.e. first a centred log-ratio transformation is applied to the count data and Principal Component Analysis (PCA) with Euclidean distance and dendrograms with Aitchison distance. The data is normalized with the Grade of Membership (GoM)^71^ analysis from the CountClust R package, therefore the raw counts from Bracken were input. ALDEx2^32^ for significant species diversity testing, includes normalization with Dirichlet distribution using 128 Monte Carlo instances, centered log-ratio transformation and significance testing with both Welch’s t-test and Wilcoxon rank test and multiple hypothesis testing with Benjamini and Hochberg false discovery rate. All plots were plotted with ggplot2^72^.

### Dental calculus protein extraction

The protein extraction was performed following the method proposed by Jersie-Christensen et al.^73^, with a blank extraction control included. To clarify the role of dental calculus in protein preservation we compared the deamidation patterns of proteins obtained from dental calculus and petrous bone of the same individuals. The sample preparation of the bone fragments closely followed that of the dental calculus. The main difference was the overnight demineralization: for dental calculus 1 ml 15-20% acetic acid was added to about 7 mg of powder while about 50 mg of the bone fragments were demineralized in 300 ml of EDTA pH 8.

After centrifugation for 10 min at 2000 g the supernatant was removed, for both sample types.

A lysis buffer (6M guanidine hydrochloride, 10mM chloroacetamide, 20mM Tris(2-carboxyethyl) phosphine Hydrochloride in 100mM TRIS pH 8.5) was then added to the powder and the pH was adjusted to 7-9. The pellet was crushed by disposable sterile micro-pestles and then incubated either at 99 °C for 10 mins (calculus) or at 80 °C for 2 hours (bone) at 500 rpm. The protein concentration was then measured by Bradford Assay.

After adjustment of pH (between 7-9), the samples were digested with rLysC (0.2 μg, Promega, Sweden) incubating under agitation at 37 °C for 2-4 hours. Subsequently, the samples were diluted to a final concentration of 0.6 M GuHCl solution adding 25 mM Tris in 10% acetonitrile. This was followed by digestion by trypsin (0.8 μg, Promega, Sweden) and incubation overnight at 37 °C under agitation.

To stop the digestion, the samples were acidified (pH <2) using 10% trifluoroacetic acid, then the proteins were collected in home-made C18 StageTips and stored in the freezer until mass spectrometry analysis.

### LC-MS

Dental calculus samples were eluted from the stage tips using 20 μL 40% ACN in water and then 10 μL 60% ACN while the bone samples by 30 μL 40% ACN in water both into a 96 well MS plate. Samples were placed in a SpeedVac™ Concentrator (Thermo Fisher Scientific, Denmark) vacuum centrifuge at 40°C until approximately 3 μL of the solution was left and then 5 μL of 0.1% TFA, 5% ACN was added.

Samples were then separated on a 15 cm column (75 μm inner diameter) in-house laser pulled and packed with 1.9 μm C18 beads (Dr. Maisch, Germany) on an EASY-nLC 1200 (Proxeon, Odense, Denmark) connected to a Q-Exactive HF (Thermo Scientific, Bremen, Germany) on a 77 min gradient. 5 μL of sample was injected. The column temperature was maintained at 40°C using an integrated column oven. Buffer A was milliQ water. The peptides were separated with increasing buffer B (80% ACN and 0.1% formic acid), going from 5% to 30% in 50 min, 30% to 45% in 10 min, 45% to 80% in 2 min, held at 80% for 5 min before dropping back down to 5% in 5 min and held for 5 min. The flow rate was 250 nL/min. A wash-blank method using 0.1% TFA, 5% ACN was run in between each sample to hinder cross-contamination.

The Q-Exactive HF was operated in data-dependent top 12 mode (dental calculus) and top 10 mode (bones). Spray voltage was 2 kV, S-lens RF level at 50, and heated capillary at 275°C. Full scan mass spectra were recorded at a resolution of 120,000 at m/z 200 over the m/z range 350–1400 with a target value of 3e^6^ and a maximum injection time of 25 ms. HCD-generated product ions were recorded with a maximum ion injection time set to 45 ms (dental calculus) and 118 ms (bones) with a target value set to 2e^5^ and recorded at a resolution of 30,000 (dental calculus) and of 60,000 (bones). The normalised collision energy was set at 28% and the isolation window was 1.2 m/z with the dynamic exclusion set to 20 s.

### Protein data analysis

Data analysis was performed with MaxQuant version 1.5.3.30^74^ with oxidation (M), Acetyl (protein N-term), deamidation (NQ), and hydroxyproline set as a variable modification and carbamidomethyl (C) as a fixed modification. Digestion enzyme was set to trypsin with a maximum of two missed cleavages allowed. For the identification, a minimum score of modified and unmodified peptides of 40 was used and a Peptide Spectral Match (PSM) and Protein false discovery rate (FDR) of 0.01 cut-off was set. All other parameters were left for the default for orbitrap mass spectrometers. Different databases and several searches were performed on the two biological matrices. First, for dental calculus, the entire SwissProt database (downloaded in January 2017) was used for the first screening and then a search against a FASTA file built using all the proteomes of the species identified by the first search and the human reference proteome from UniProt (downloaded in August 2018) was performed. Likewise, the bone fragments were searched just against the human reference proteome from UniProt. The peptides identified were then filtered applying quality controls. First of all, the resulting proteins group output was filtered to remove reverse and common contaminants. Furthermore, all protein groups with a value of Razor + unique less than two were removed. The spectra for each peptide associated with a proteins group were then manually validated and the species was confirmed by BLAST search^75^.

Finally, we calculated the asparagine and glutamine deamidation rate using the python tool proposed by Mackie et al.^33^ in order to assess molecular damage associated with the antiquity of the remains. The deamidation rate was evaluated on all reported proteins in Table 1 and also separately on collagen proteins identified in dental calculus and bone samples (SI Appendix).

## Acknowledgments

We would like to thank Eske Willerslev for financial support. This project has received funding from the European Union’s Horizon 2020 research and innovation programme under the Marie Sklodowska-Curie grant agreement No 751349 allotted to G.S. G.S. also received support from the SYNTHESYS Project http://www.synthesys.info/ which is financed by European Community Research Infrastructure Action under the FP7 “Capacities” Program.” Support for this project was also provided by PRIN MIUR (Italian Ministry for the Universities) 2009-11 (3 years) prot.2010EL8TXP National Scientific Coordinator and Principal Investigator OR: Biological and cultural heritage of the central-southern Italian population through 30 thousand Years EPIC. The Novo Nordisk Foundation Center for Protein Research (CPR) is funded in part by the Novo Nordisk Foundation (Grant number NNF14CC0001). This work was also supported by the Lundbeck Foundation, the Novo Nordisk Foundation and the Wellcome Trust (grant no. WT104125MA). We would like to thank Benedetto Sala for his suggestions in order to identify wild animal species inferred by proteomic dental calculus data.

## Author contributions

M.S., E.C, M.E.A., O.R. and G.S. initiated the project on Multi-omics analysis. G.S., A.M., C.M.L., D.L.V., F.M. and P.F.F. collected the samples. G.S. performed the human genetic extraction and library preparation, G.S. and M.M. performed the proteomic extraction on dental calculus, M.M. carried out mass spectrometry runs, with support and resources provided by J.V.O. A.K.F. performed the genetic extraction from dental calculus and library preparation, S.H.F. carried out the dental calculus metagenomic data analysis, G.S. carried out human genetic and proteomic dental calculus data analysis. M.C. provided palaeobotanical input for plants consumption interpretation inferred by proteomic data. D.L.V and F.M. provided input about the archaeological context and the radiocarbon date published in the present paper used for the contextualization and interpretation of data obtained. P.F.F. carried out the human bones morphological analysis. M.S. and E.C. provided supervision of data analysis and supervised the interpretation of the results and the formulation of the conclusions. G.S., M.S. and S.H.F. wrote the manuscript and all authors reviewed and approved it. D.L.V. wrote the archaeological part of the manuscript. P.F.F. wrote the anthropological part of the manuscript. The data published fall into an overall project about the revision of the archaeological and anthropological collections from San Teodoro curated at the Museo e Istituto Fiorentino di Preistoria (Florence) designed by F.M., D.L.V. and P.F.F.

