## Supplementary Information for "Genomic ancestry, diet and microbiomes of Upper Palaeolithic hunter-gatherers from San Teodoro cave (Sicily, Italy)"

### Introduction of the archaeological site of San Teodoro: dating and anthropological analysis

*Geographical setting* **(D.L.V.)**

The San Teodoro site near Acquedolci (Messina, North-Eastern Sicily), is a large cave (about 60 m long, 20 m wide and up to 20 m high) which opens in the Jurassic limestone of Pizzo Castellaro, at the toe of the northern side of Monte S. Fratello (Nebrodi Mountains). The cave is located at an altitude of 135 m, about 2 km south from the Tyrrhenian coast. The site stands between two rivers: Inganno, about 2 Km Est, and Furiano, about 2,5 km West (Fig. S2).

*History of researches and Archaeological setting* **(D.L.V.)**

After its discovery by Baron Anca in 1859, several scholars were interested in the prehistoric remains of San Teodoro cave. The first human remains were recovered in 1937 by G. Bonafede who unearthed the burial San Teodoro 1[1]. Other skeletal remains related to 3 distinct individuals were recovered a few years later by Maviglia[1]. In 1942 Graziosi conducted first systematic archaeological excavations[2], where he detected a detailed stratigraphic sequence and recovered three other individuals.

New modern researches were carried on from 1998 to 2006 by Bonfiglio who conducted several excavations in the Pleistocenic deposit, bearing endemic mammal fauna, underlying the anthropogenic one.

The archaeological sequence detected by Graziosi consists in two main levels, a lower level, subdivided into three non-anthropogenic layers (E-F), rich in faunal remains (among which the endemic *Elephas mnaidriensis* and the hyena *Crocuta crocuta spelaea*), and an upper level, subdivided into four anthropogenic layers (A-D) containing abundant lithic artefacts related to the Late Upper Palaeolithic (Late Epigravettian) and several faunal remains: red deer (*Cervus elaphus*), which remains are prevalent in the faunal assemblage, wild boar (*Sus scrofa*), abundant, aurochs (*Bos primigenius*), scarce, wild ass (*Equus hydruntinus*), wolf (*Canis lupus*), fox (*Vulpes vulpes*) hyena (*Crocuta crocuta spelaea*) very scarce. The latter two species occur only in the lower part of the anthropogenic deposit (layer D). The few remains of hyena could be intrusive in layer D because of the burial pits dug by the epigravettians in the underlying non-anthropogenic deposit. In the whole anthropogenic sequence few mollusc shells (both marine and terrestrial) occur. The human Upper Palaeolithic skeletal sample from San Teodoro (ST) is presently composed by at least seven individuals (Table S1), as several other fragmentary human remains have not yet been fully analyzed.

Four individuals (ST 1-4) were found in layer E, below a red ochre lens, but they pertained to the beginning of the cave occupation related to layer D[2]. Individual ST 5 was found over the red ochre lens, nevertheless its absolute chronology is consistent to the other dated human remains (ST 1 and ST 4; see Fig. S3 and Table S2); probably it is the result of an *ab antiquo* displacement related to a disturbance of a burial of which the original position was in the same layer as the others.

Regarding the chrono-cultural framework, stone tool assemblages from layer A-D show the typical traits of the local Final Epigravettian industries. Recent techno-typological studies suggest an attribution of San Teodoro lithic industries to a later stage of this culture in Sicily, chronologically referred to around 12-11.000 uncal. BP. Considering that layers A-D are imputable to human frequentations which are subsequent to the burials standing in the layer E, the chronology based on techno-typological features of lithics is consistent with both their stratigraphic position and AMS radiocarbon dates from ST 1[2], ST 4 and ST 5 individuals (Table S2). Here, we also present the new radiocarbon date of the individuals: ST 4 and ST 5 (Table S2).

ST 4 and ST 5 refer to the same period, spanning from about 15,300-14,200 cal. BP. We also tried to date ST 3, however the dating was unsuccessful. The chronology of ST 4 and ST 5 match with the AMS measure of ST 1[3] (Table S2 and Fig. S3). The AMS date of ST 5, coming from layer B, is consistent, although a little older, with the chronology of the other individuals and leaves open the issue of the original position of these human remains, which Graziosi[4] hypothesized could be in a secondary position.

<u>Anthropological analysis</u> **(P.F.F.)**

Graziosi[2], in his original publication, attributed to the male sex five of the Upper Palaeolithic individuals from San Teodoro cave (ST 1-5). Two other individuals were published in the following years, ST 6, Pardini[4], and ST 7, Aimar and Giacobini[5]. These two individuals are represented by crania only, the former is a fragmentary frontal and facial skeleton, the latter is lacking the right emi-frontal and the facial skeleton. ST 6 has been sexed as female and ST 7 as male. Later, one of the current authors, Fabbri[6], proposed that ST 1 and ST 4 should be diagnosed as females on the basis of pelvis morphology. As to general cranial morphology, compared to Late Upper Palaeolithic individuals, ST 3 shows very large and robust mastoids and pronounced supraorbital and nuchal reliefs while ST 5 is more gracile in the three features (Fig. S4).

The female sex determination for ST 6 seems reasonable as this individual is certainly the most gracile among the seven known crania from ST while it seems likely that the robust ST 7 is a male. ST 2 has intermediate cranial robustness and no other bones are preserved; sex determination should be viewed as impossible on morphological grounds.

The sex determination of ST 3 and 5 among the samples from San Teodoro was evaluated by some dental, cranial and postcranial measures that can be taken in at least 4 of the seven individuals: maximum cranial length (Martin M1); occlusal upper canine area (MD*BL); humeral lower epiphysis breadth (Martin M4). These measures are known to show a higher sexual dimorphism in modern humans, and they have been compared to those recorded in pelvis sexed Upper Paleolithic Italian individuals: Barma Grande (BGR), Romito (ROM), Vado all'Arancio (VAR), Villabruna (VIL)[7] and Arene Candide (ACA)[8].

Maximum cranial length (M1) can be measured in ST 1, 2, 3, 5 and 7 (Fig. S5). This measure doesn't seem to have a high sexual discriminatory power: pelvis sexed gracile female ROM 1 and robust male VIL have nearly identical measures, respectively 180 and 181 mm; pelvis sexed male ACA 4 and female ROM 4, are both 195 mm. Maximum measure in ST samples (198 mm) is recorded in pelvis sexed female ST 1, whose measure is identical to the ones recorded in pelvis sexed males ROM 7 and 8 and very close to ST 3 (196 mm) and ROM 4 (195 mm). ST 7 very robust cranium is shorter than more gracile ST 5 cranium, respectively 187 and 192 mm.

As to occlusal upper canine area (Fig. S6), it could be computed in individuals ST 1, 2, 3 and 6, all of them are placed in the lower half of Italian UP range and ST 6 shows the lowest value. Pelvis female sexed ST 1's value (68.03 $mm^2$) is slightly larger than the one observed in ST 3 (63.04 $mm^2$) and both of them are lower than unsexed ST 2 (70.94 $mm^2$). As observed for maximum cranial length, occlusal upper canine area doesn't permit clear discrimination between sexes: values computed for the four individuals from ST are lower than those observed in female pelvis sexed BGR 3 and ROM 4.

The only postcranial bone measurement recordable in both ST 3 and ST 5, as well as in pelvis sexed individuals ST 1 and ST 4, is humeral lower epiphysis breadth (Table S3).

The four individuals from San Teodoro show limited metrical variation. The three females, ST 1, 4 and 5, are very similar, respectively 60, 57.5 and 59 mm, and fall in the range 52-60 mm where male and female variabilities overlap, the only ST 3 sample (62 mm) is slightly over the female maximum.

The three chosen measures recorded in ST samples give conflicting results when compared to other Upper Palaeolithic Italian samples. In ST samples, canine occlusal areas are generally small, maximum cranial length spans most of Italian Upper Palaeolithic variability, and humeral lower epiphysis breadth are mostly in the overlapping area of male and female ranges, but we cannot exclude that this is at least partially due to the small size of comparison samples. These observations confirm that without diagnostic pelvic features, sex determination based on available singular cranial, dental or postcranial measures are not reliable except when dealing with individuals placed at the upper edge of male range or at the lower one of female range.

Cranial morphological features commonly used for sex determination, supraorbital and occipital reliefs and mastoid size are scored following Walker[9] (Table S4). ST 3 is one of the more robust crania in Upper Paleolithic Italy and the most robust among San Teodoro individuals while ST 5 shows robust supraorbital reliefs (score 4) and gracile mastoid and nuchal crest (score 2 for both).

Considering all the information obtained from ST samples, we find more women (ST 1, 4, 5, 6) than males (ST 3, 7), and ST 2 is not sexable on the basis of cranial measures and features.

### Deamidation pattern comparison between petrous bones and dental calculus

Furthermore, we compared the deamidation patterns of the peptides obtained from the dental calculus and from the petrous bones of the same individuals, in order to verify and confirm different preservation states between the two matrices analyzed. The peculiar composition of dental calculus allows it to protect trapped biomolecules from environmental attack[10], unlike bone, where endogenous biomolecules are more easily subjected to environmental factors. By comparing the collagen deamidation patterns obtained on bone and dental calculus, San Teodoro 3 (Fig. S33) shows an almost identical preservation state. While, the collagen deamidation rates obtained from the bone and from the dental calculus in San Teodoro 5 (Fig. S34) shows a better preservation state of the dental calculus proteins. To date, it has not yet been clarified if the number of total identified proteins and their diagenesis in dental calculus is due to the mechanism associated with dental calculus formation or the taphonomic factors to which it could have been subjected.

By comparing the damage patterns of collagen obtained in dental calculus versus bone from the same individual, we speculate that since the two individuals were recovered from two different areas within the cave, the calculus samples have been subjected to different environmental factors leading to two different protein contents and preservation states. Surely, further studies are needed to address this question.

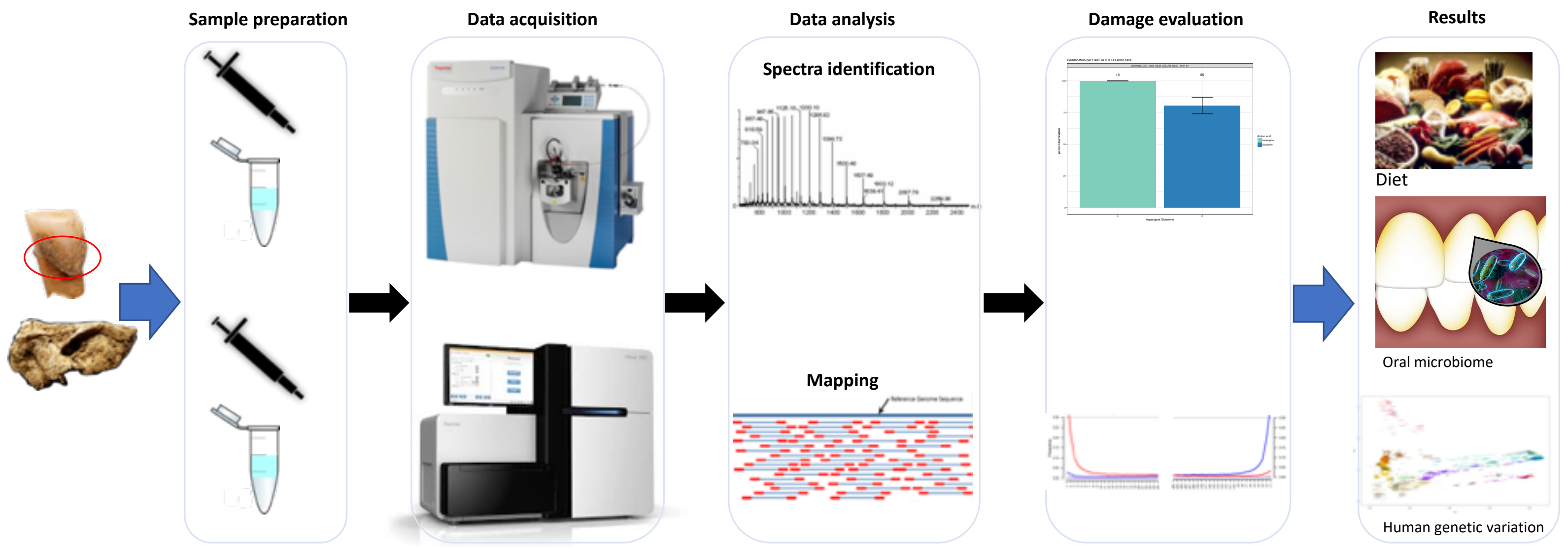


**Fig. S1:** Multiproxy approach used.

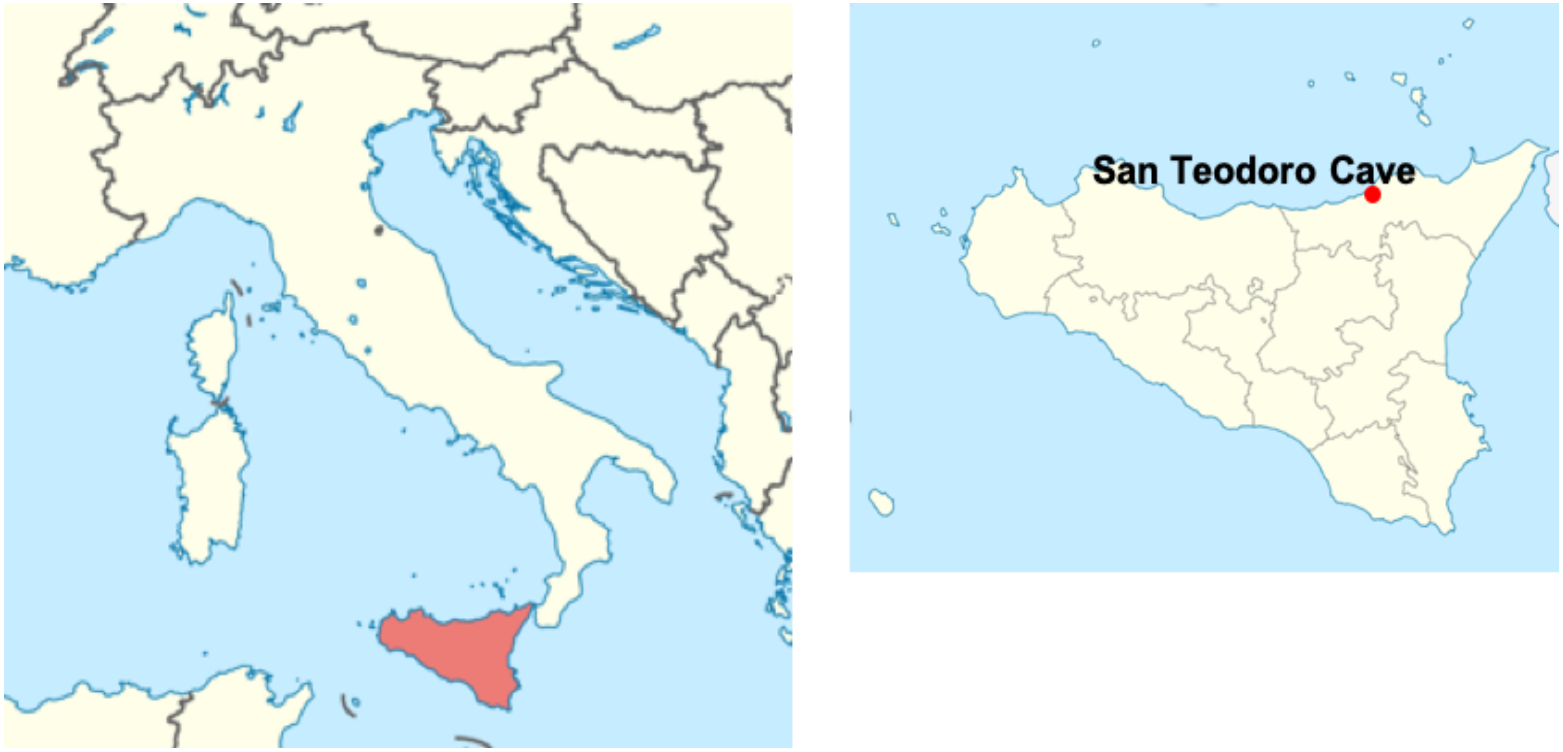


**Fig. S2:** Sampling location of San Teodoro cave (Sicily, Italy).

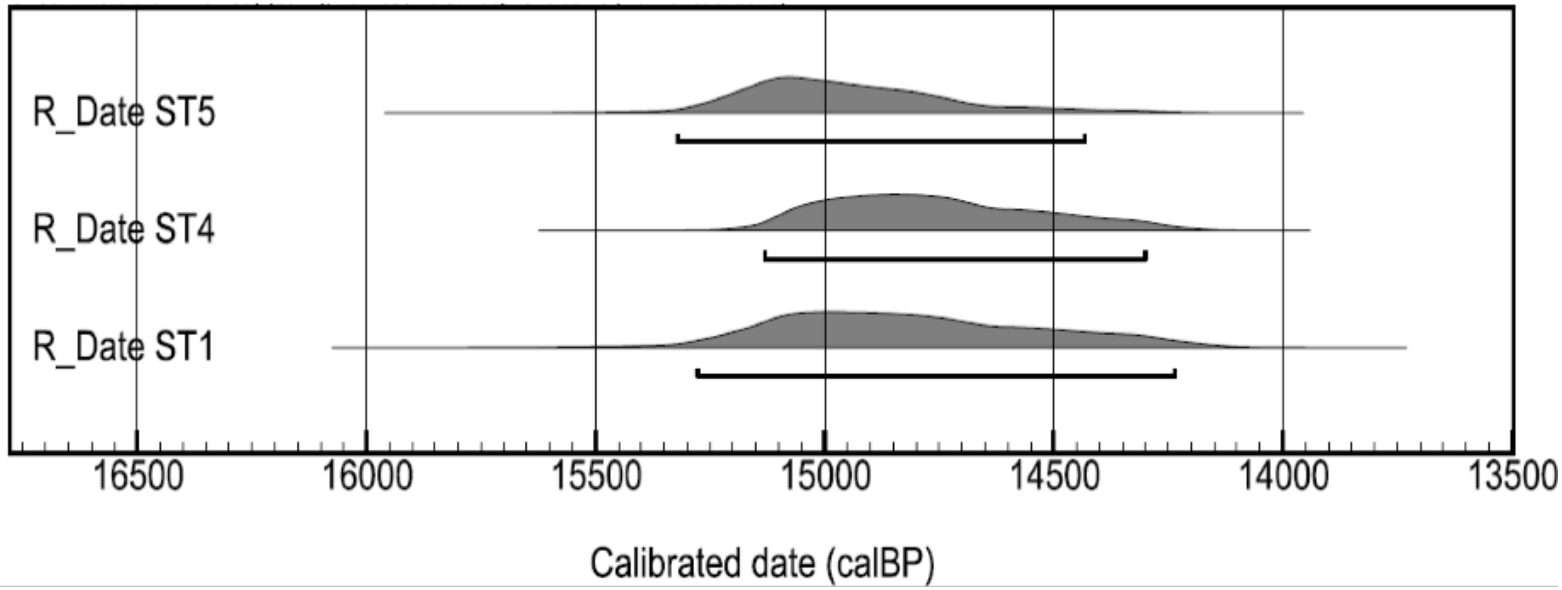


**Fig. S3:** Cumulative calibration curves of the AMS Radiocarbon dates on human samples from San Teodoro individuals ST1, ST4 and ST5.

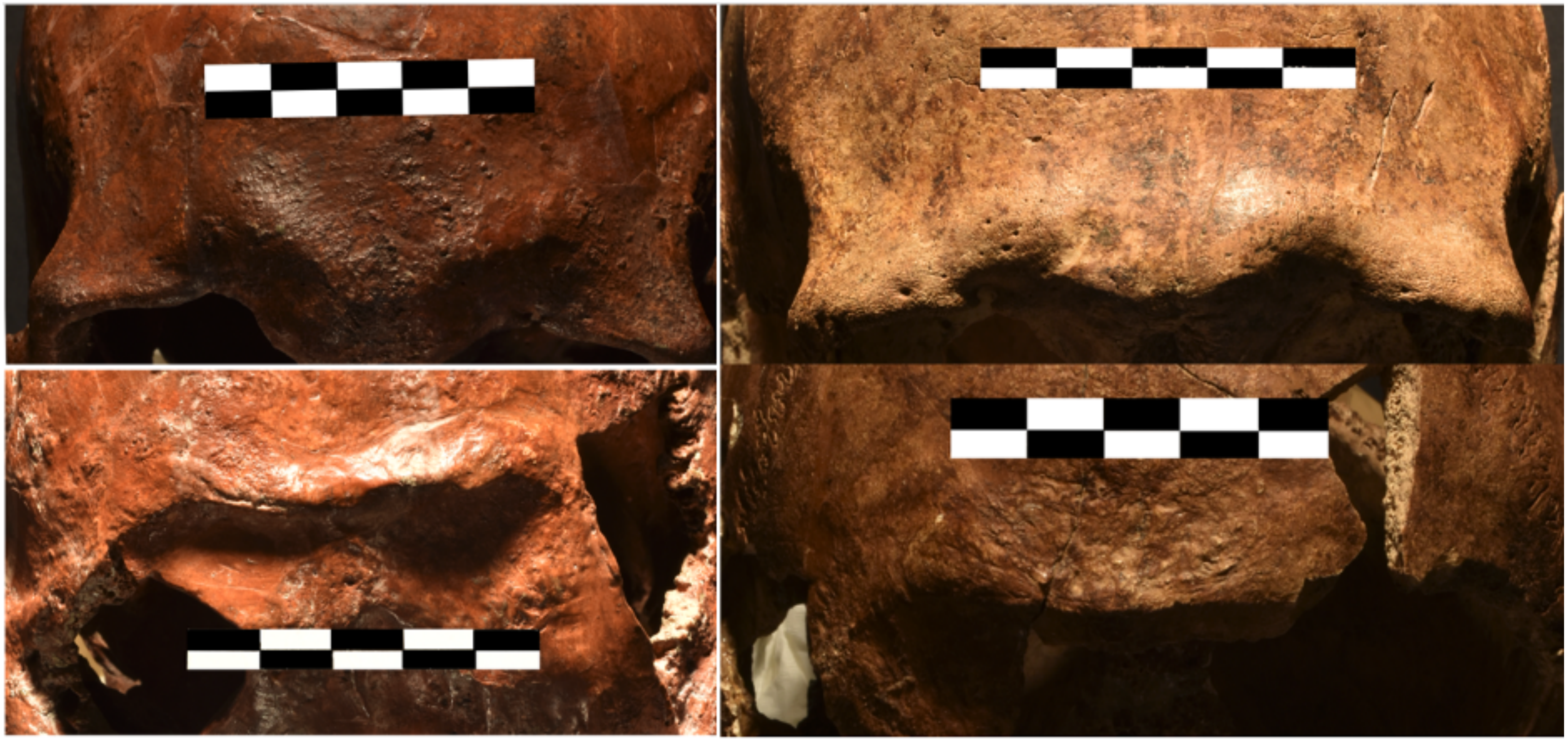

**Fig. S4:** Left San Teodoro 3 and right San Teodoro 5 skulls from top to bottom: supraorbital reliefs; occipital reliefs. Scale bar 5 cm.

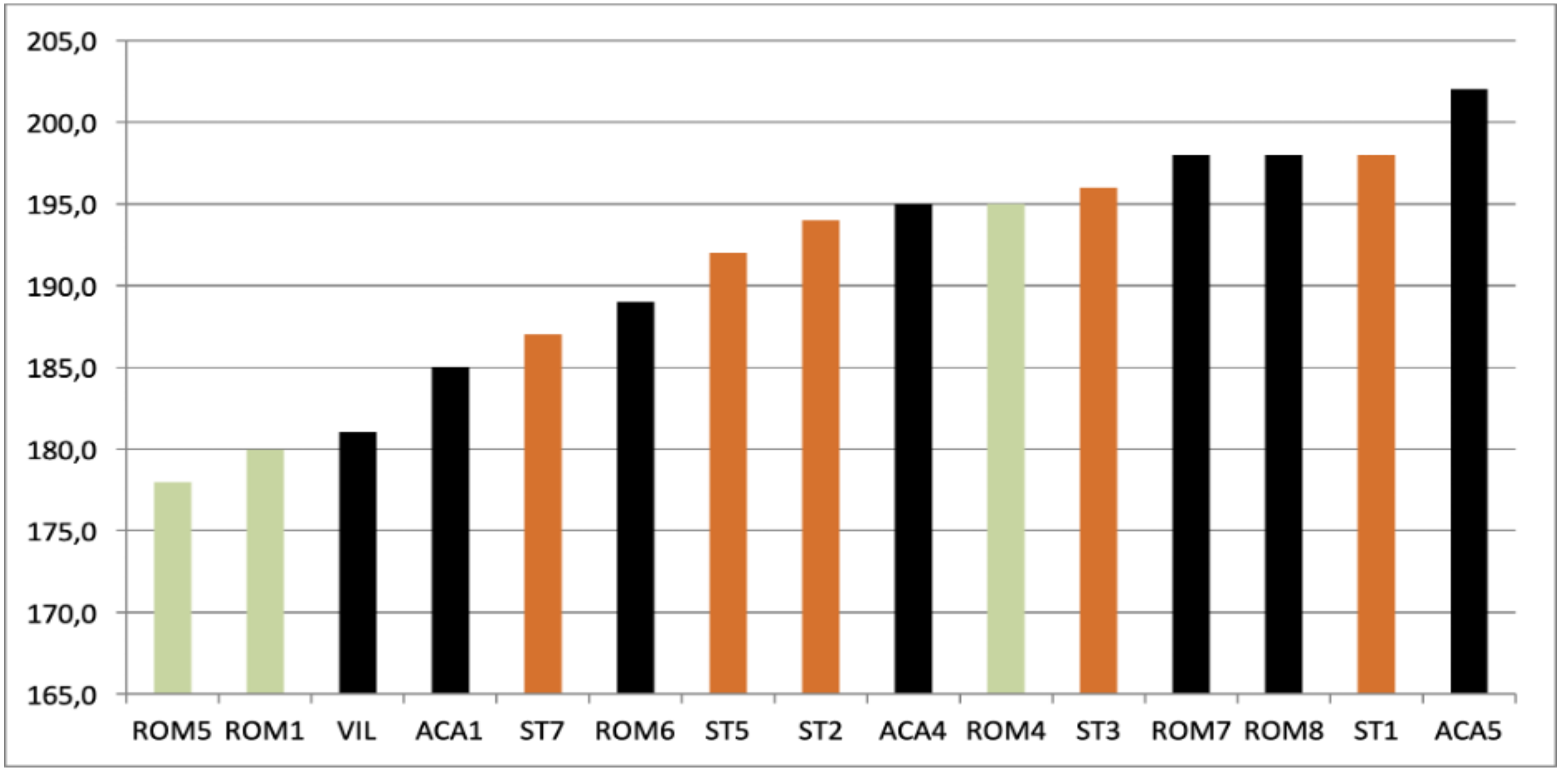


**Fig. S5:** Maximum cranial length (M1) in Italian Upper Palaeolithic upper canines. San Teodoro in orange; male in black; female in light green.

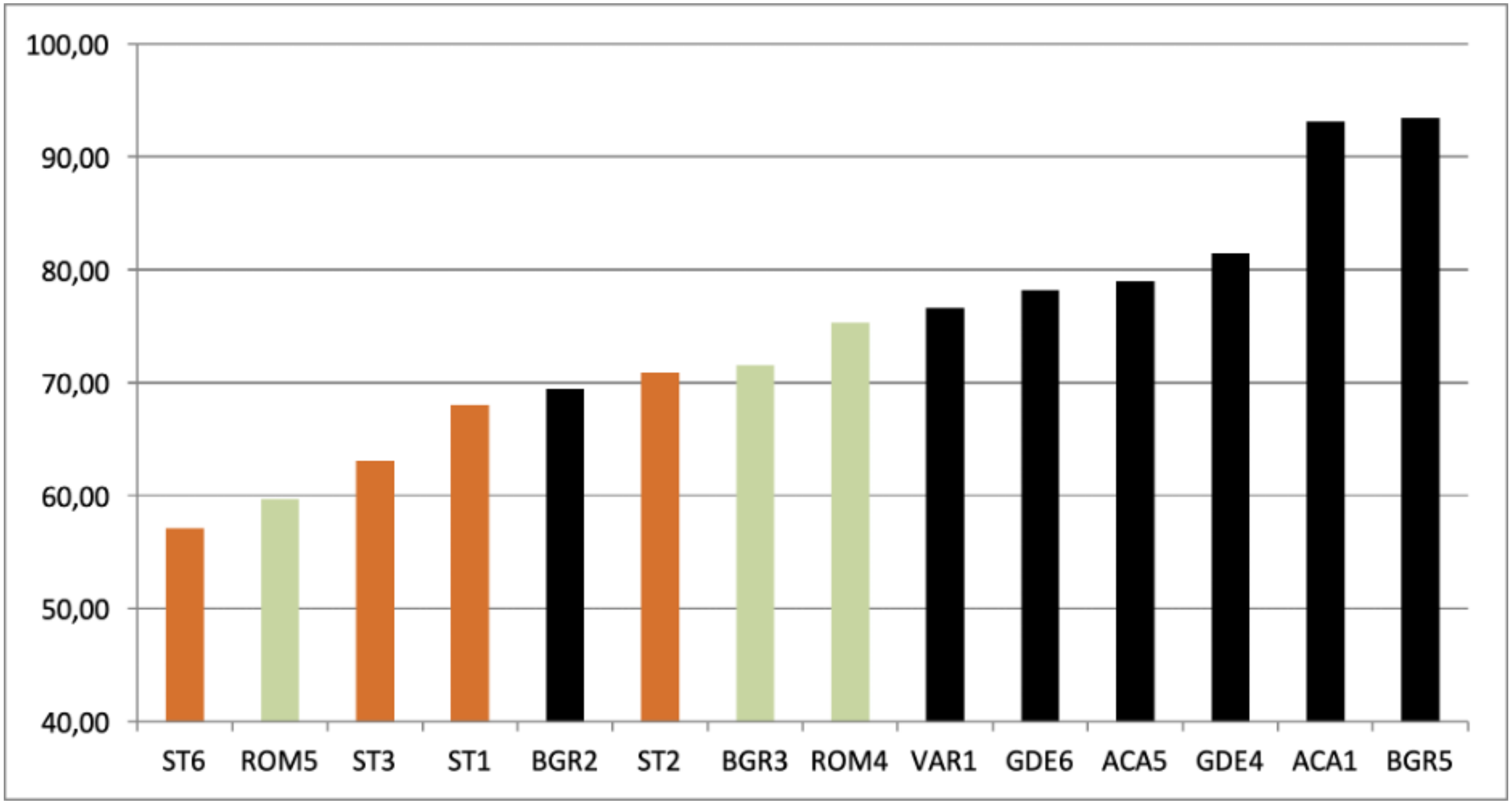


**Fig. S6:** Occlusal area (MD*BL) in Italian Upper Palaeolithic upper canines. San Teodoro in orange; male in black; female in light green.

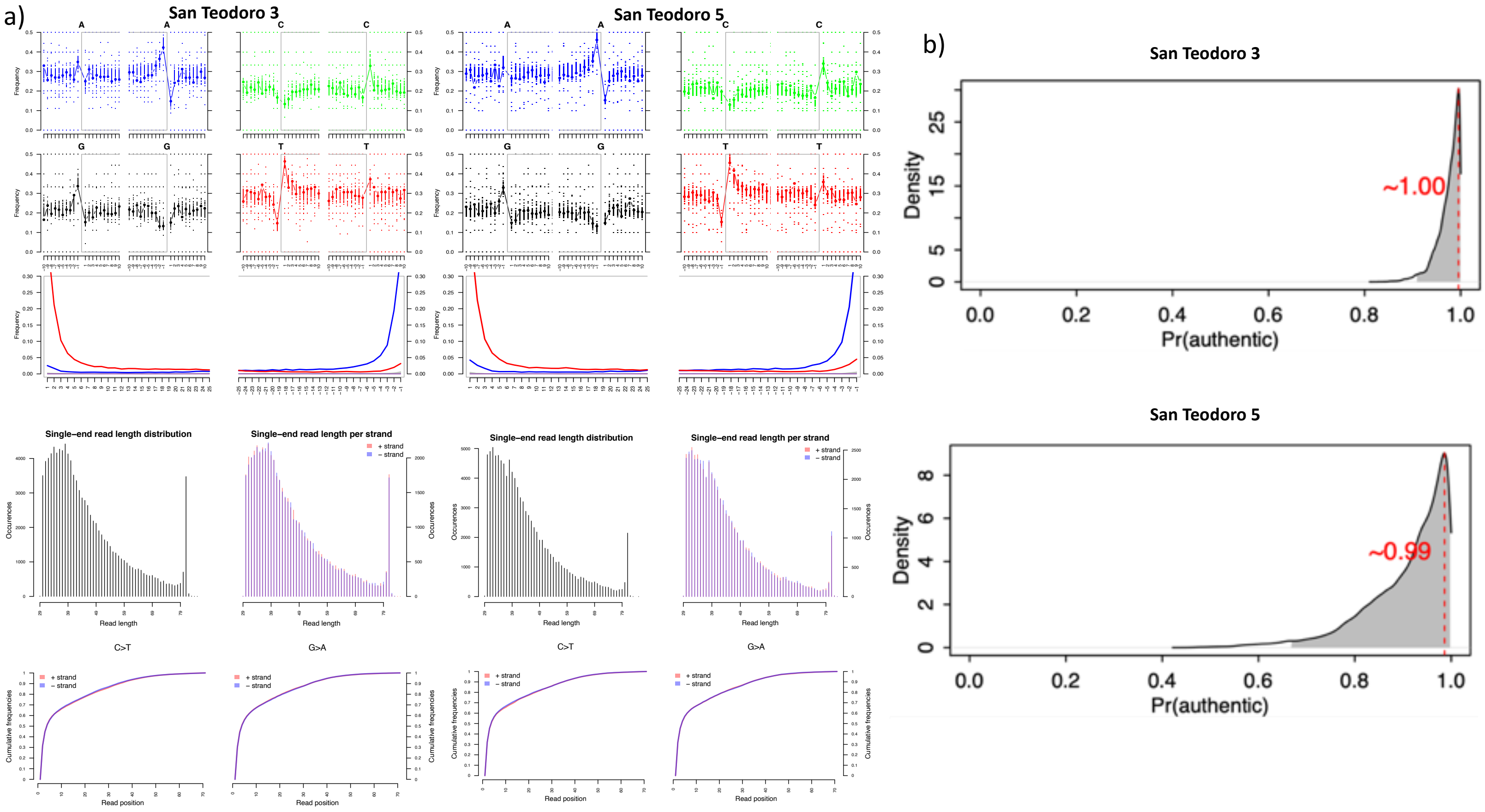


**Fig. S7:** a) Damage patterns and reads length of human DNA of samples analyzed. b) Results of Likelihood-based mitochondrial contamination estimates by contamMix.

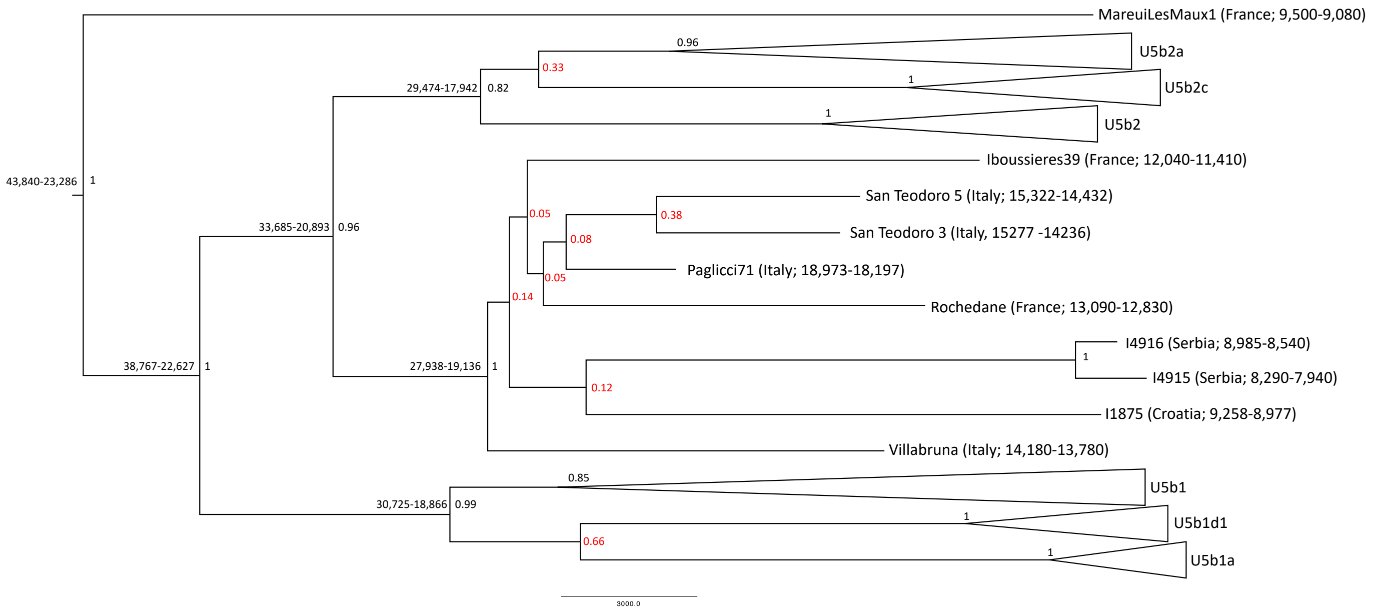


**Fig. S8:** Phylogenetic tree of U5b haplogroup based on 31 ancient samples (Table S7) and using MareuilLesMeaux1 as outgroup (U5a). The San Teodoro 3 date coming from other individuals found in the same layer (Table S2). Estimated divergence dates for principal nodes, as well as bootstrap values associated with each node are shown, with bootstrap values lower than 0.8 indicated in red.

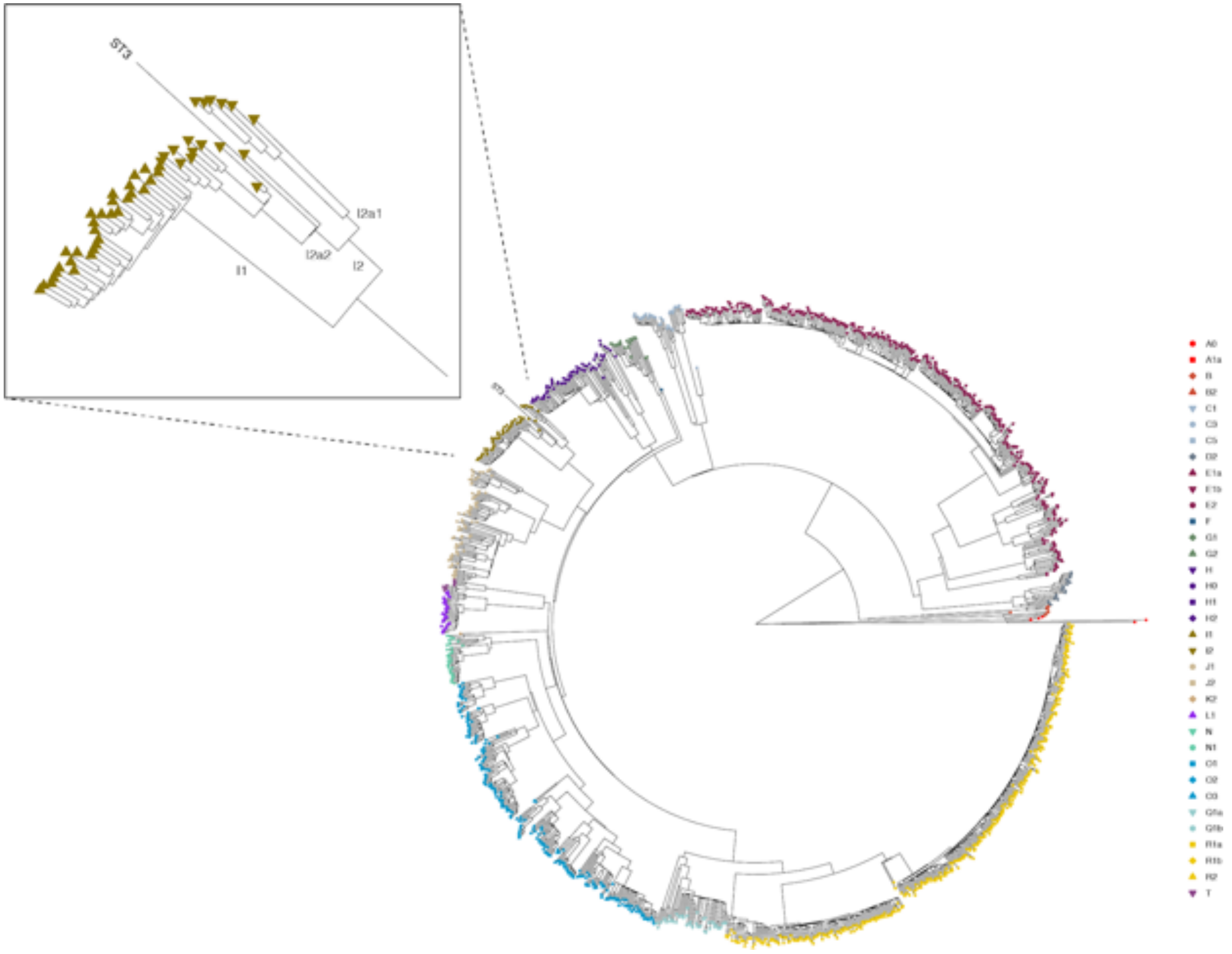


**Fig. S9:** Phylogenetic placement of San Teodoro 3 Y chromosome. Phylogenetic tree of Y chromosome sequences from the 1000 Genomes project, with maximum likelihood placement of ST3 using epa-ng. Major haplogroups are indicated with colored symbols.

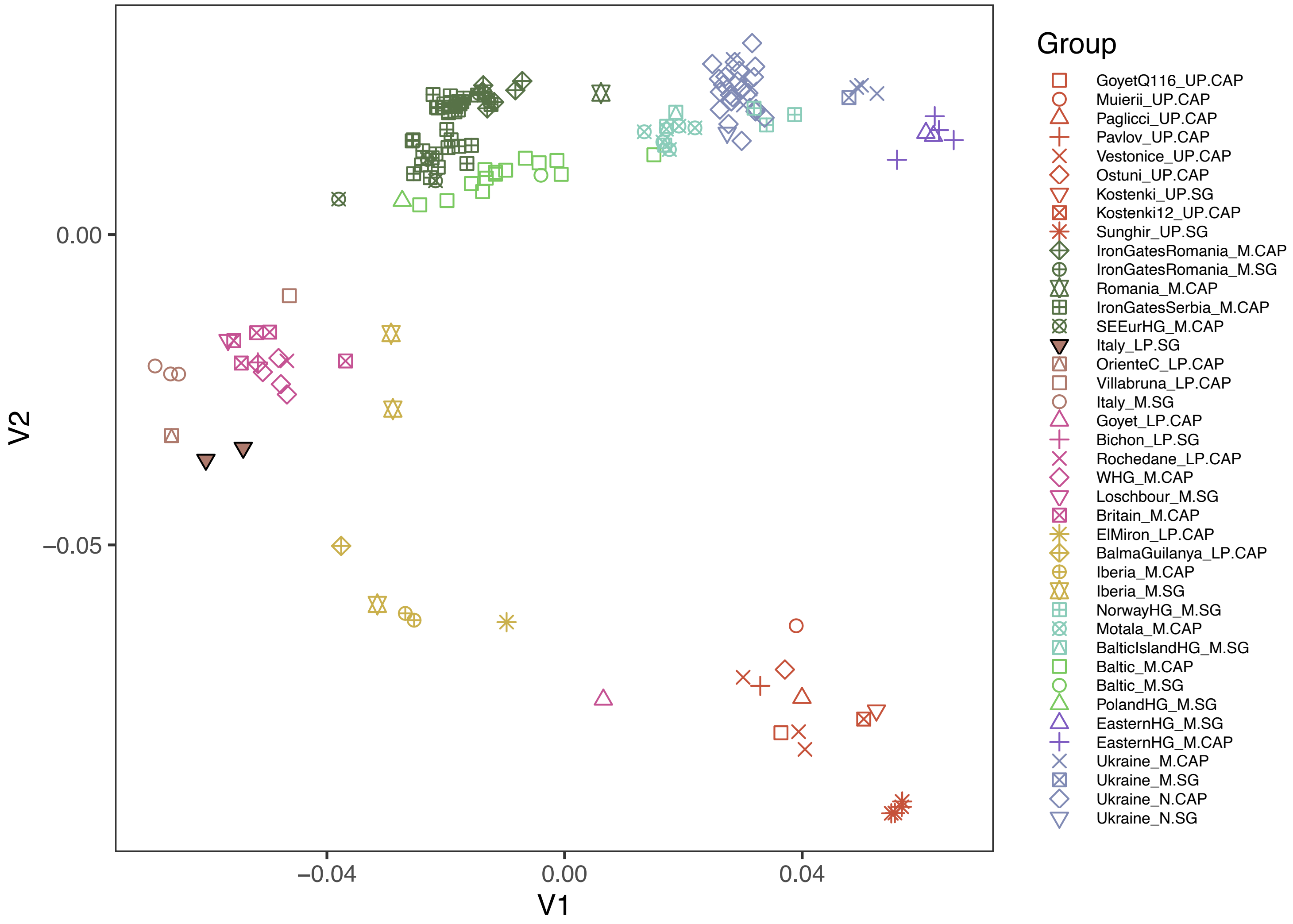


**Fig. S10:** Multidimensional scaling (MDS) of 160 pre- and post-LGM hunter-gatherer individuals, based on a pairwise identity-by-state (IBS) allele sharing.

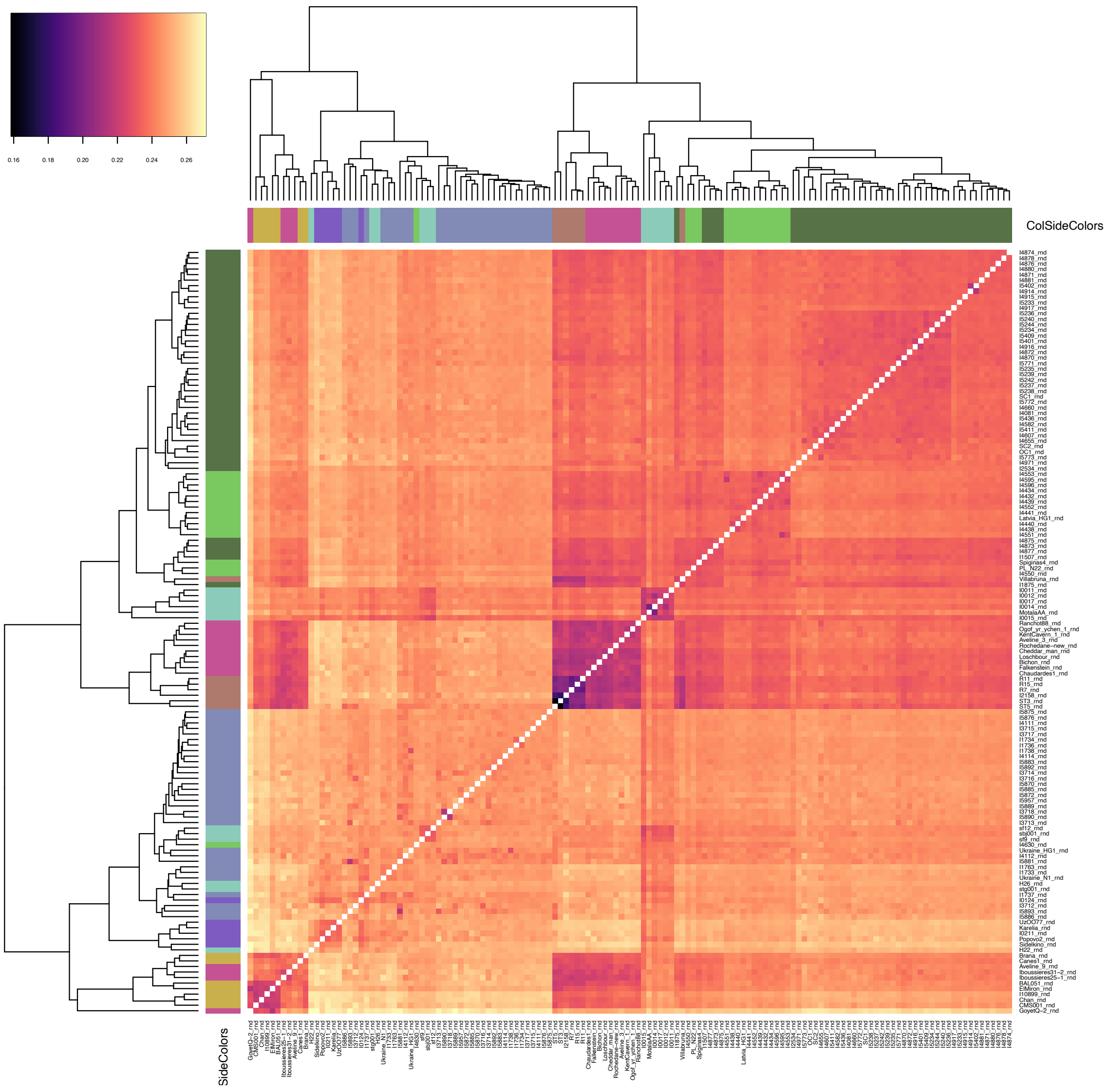


**Fig. S11:** Heatmap of pairwise genetic distances between individuals, calculated as 1- p(IBS).

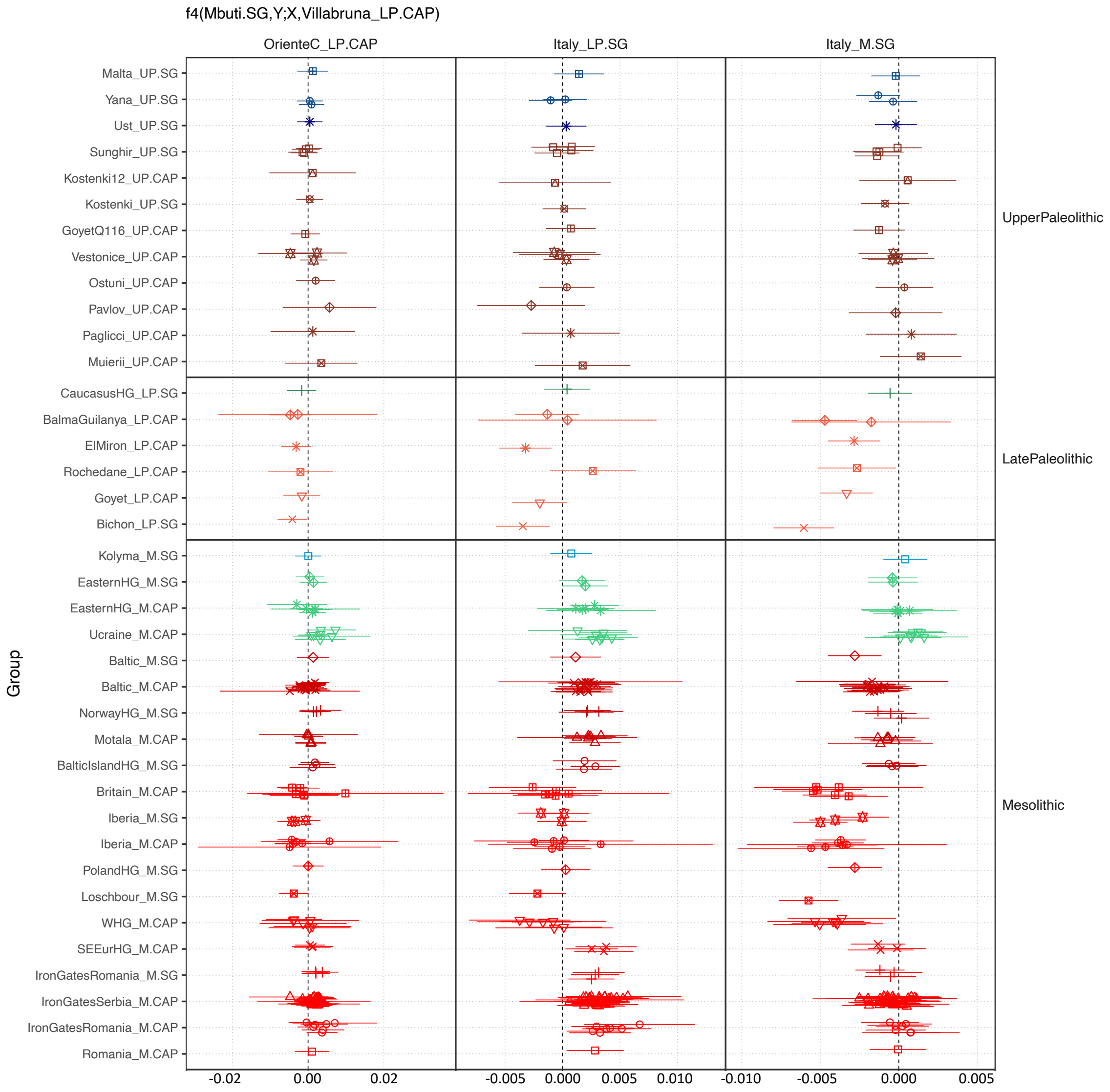


**Fig. S12:** Statistics $f_4$(Mbuti, Y; X, San_Teodoro_LP.SG).

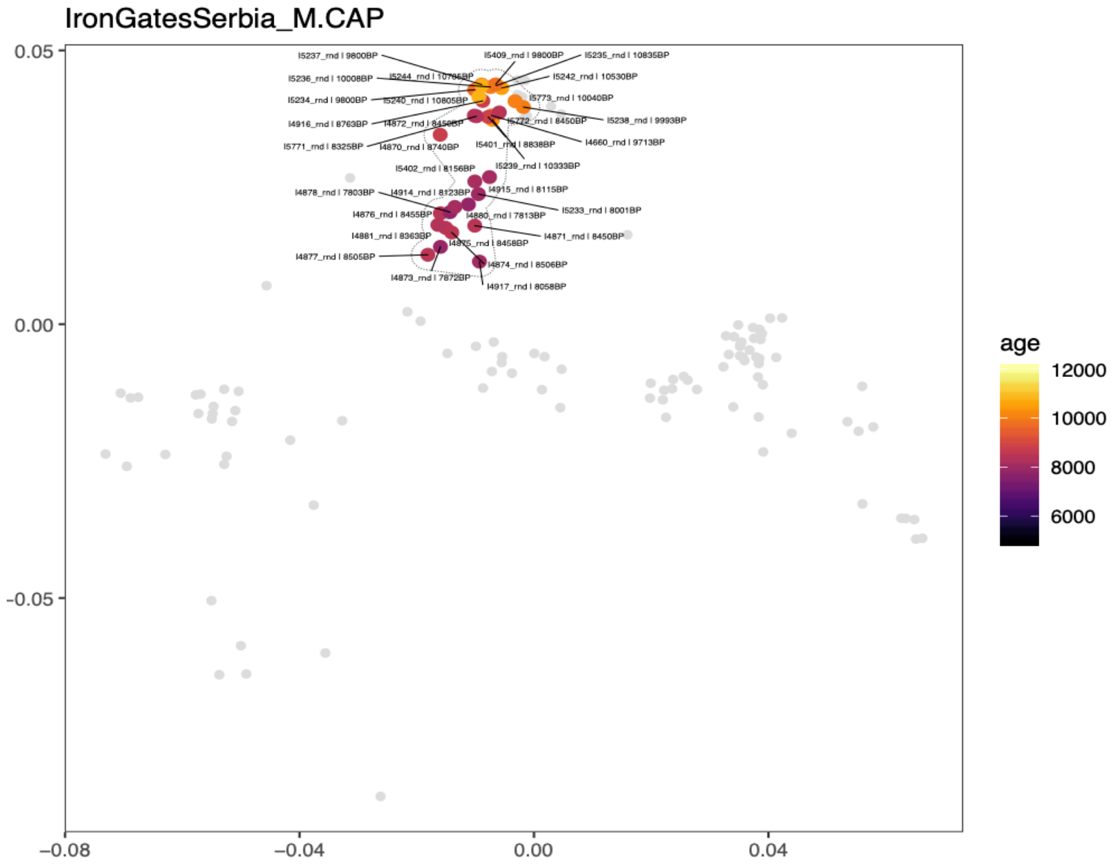


**Fig. S13:** MDS plot showing age-related stratification of hunter-gatherer individuals from Iron Gates, Serbia.

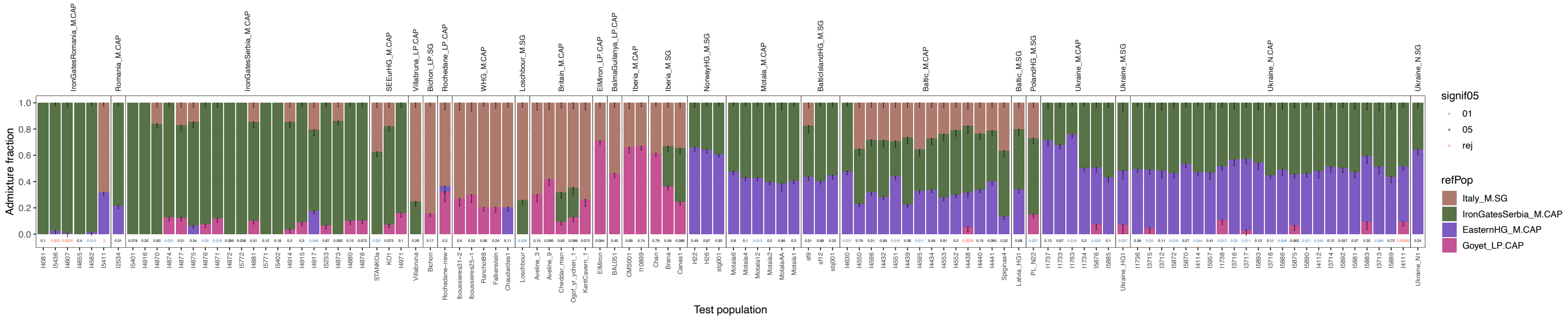


**Fig. S14:** Ancestry proportions of post-LGM Hunter-Gatherers, inferred using *qpAdm* with the associated p-value.

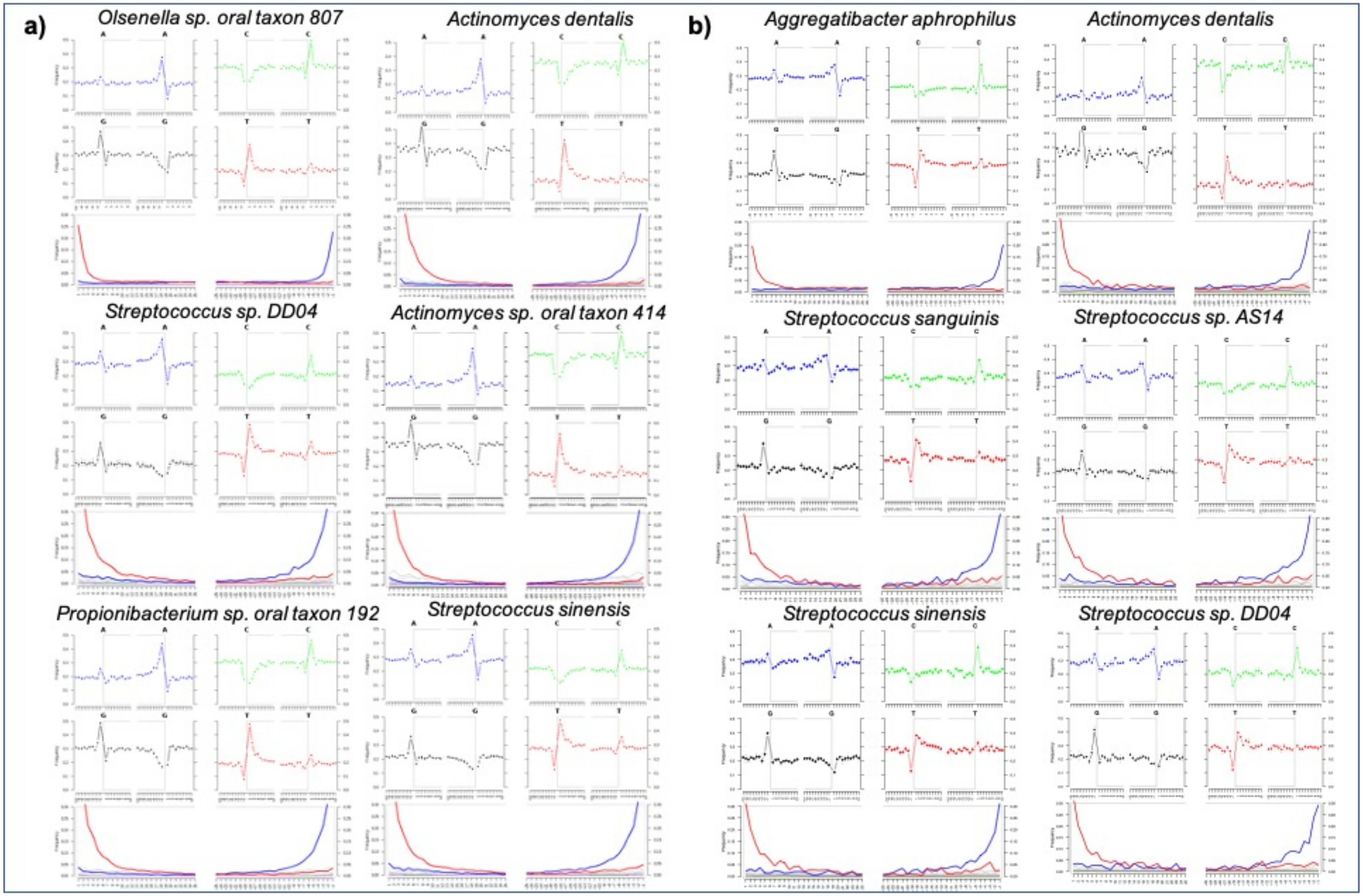


**Fig. S15:** Damage plots for the most abundant oral species identified in San Teodoro 3 (A) and in San Teodoro 5 (B) (Table S16).

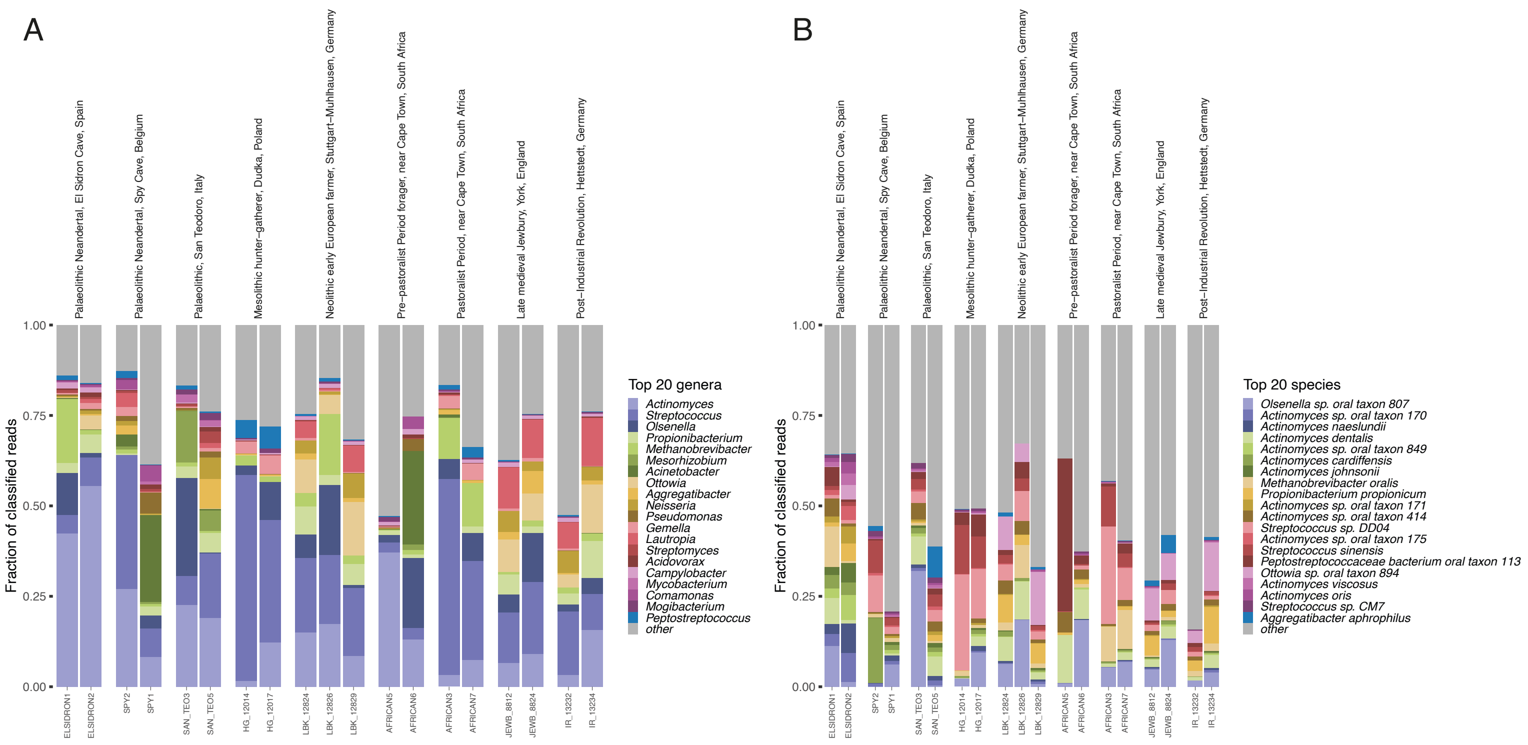


**Fig. S16:** Relative abundances of the top 20 most abundant a) genera (Table S15), b) species in the ancient calculus samples (Table S16).

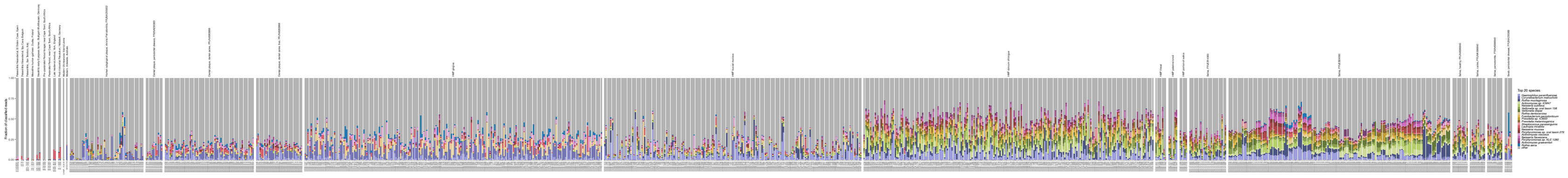

**Fig. S17:** Relative abundance bar_plot of the top 20 most abundant genera in all oral samples.

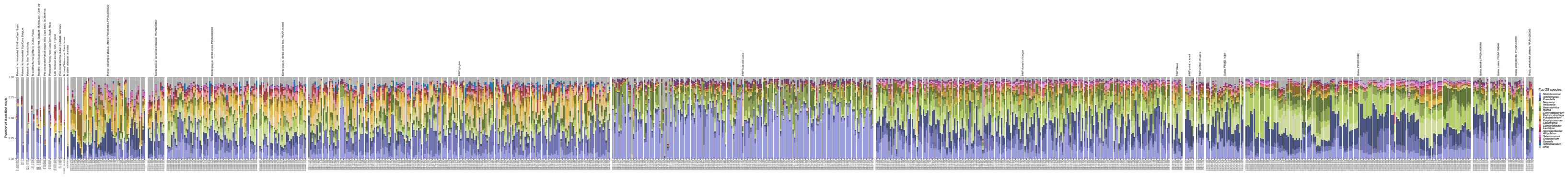

**Fig. S18:** Relative abundance barplot of the top 20 most abundant species in all oral samples.

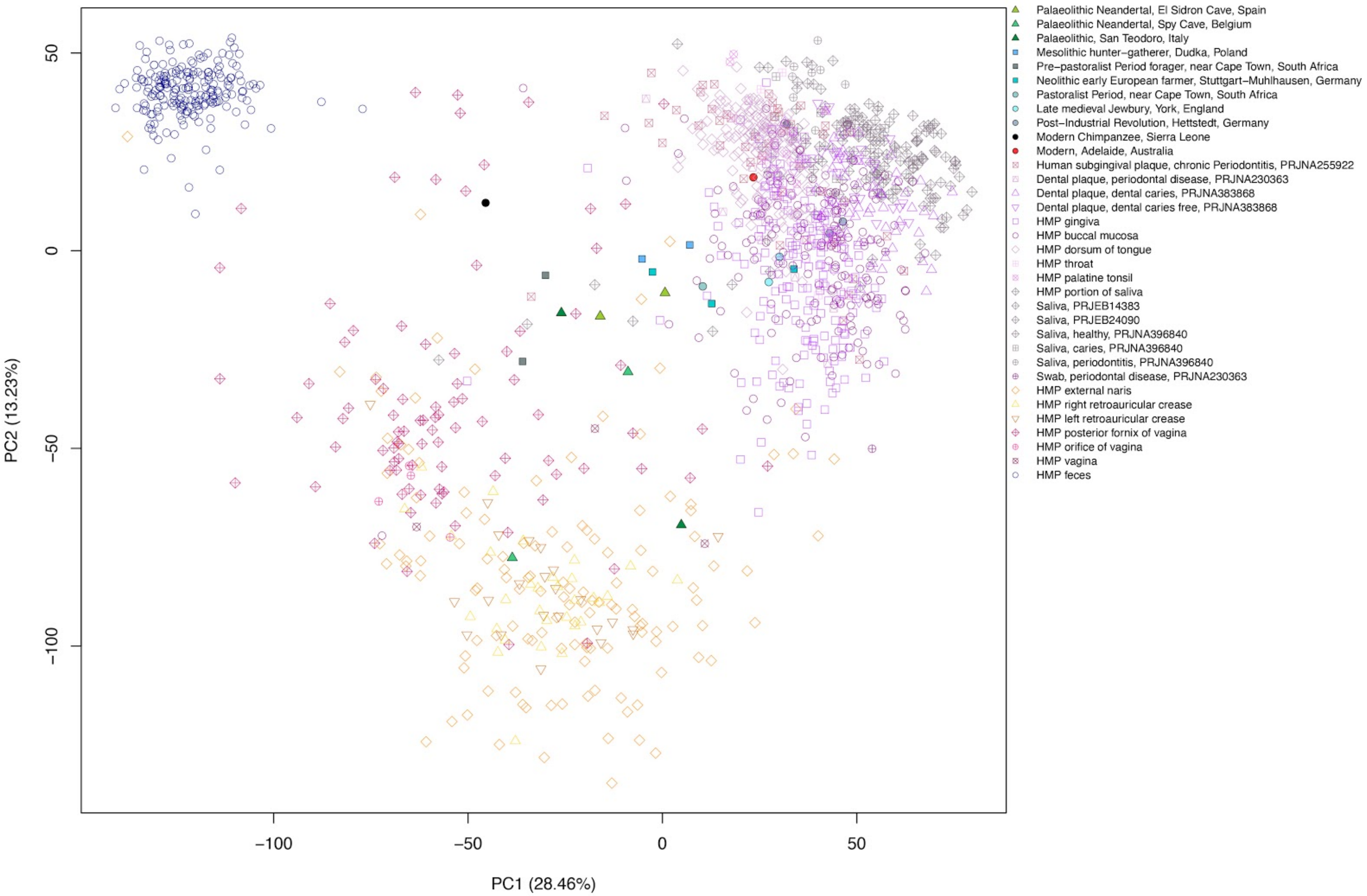


**Fig. S19:** PCA, all samples, clr transformed.

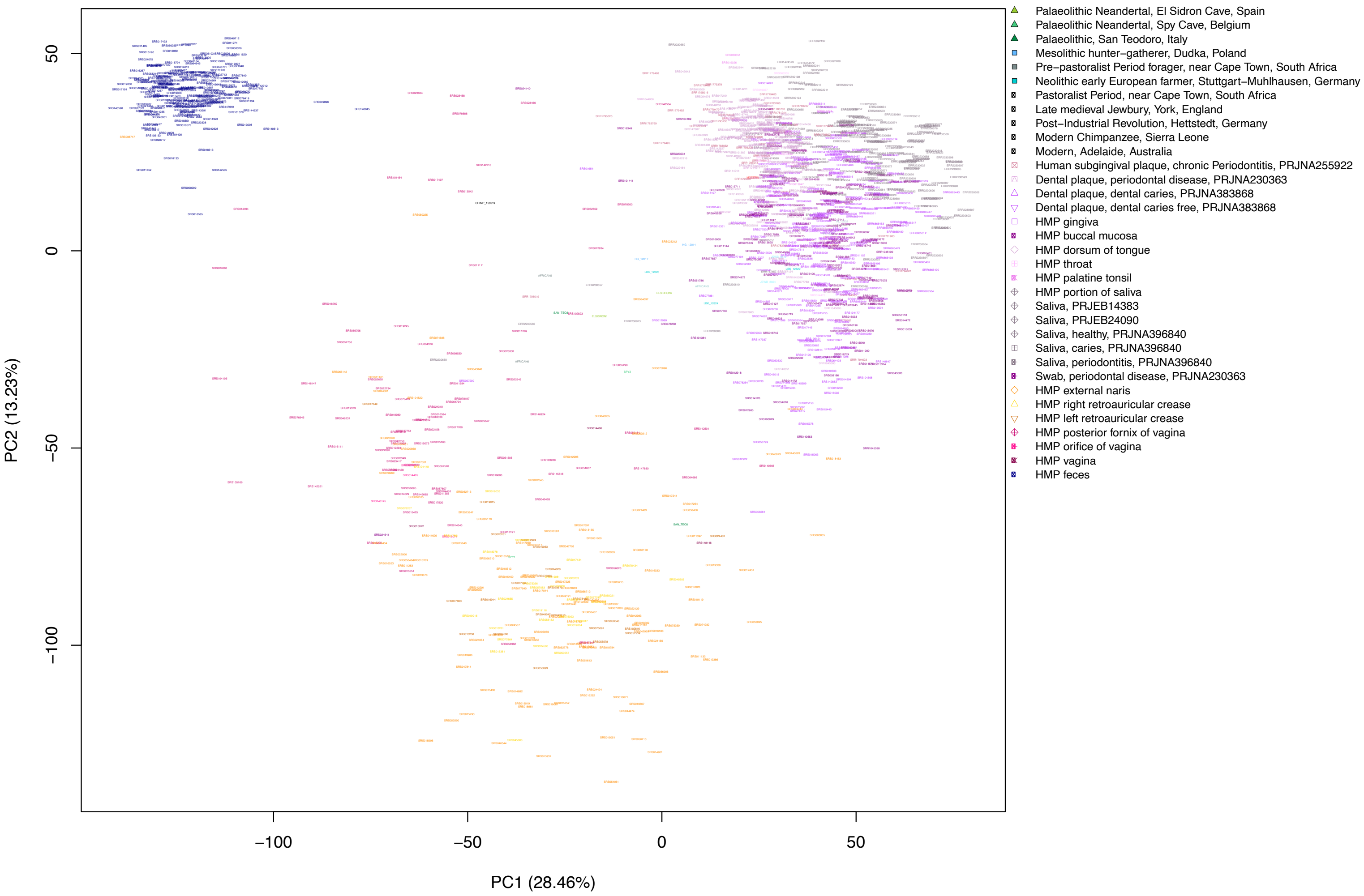


**Fig. S20:** PCA all samples, including labels.

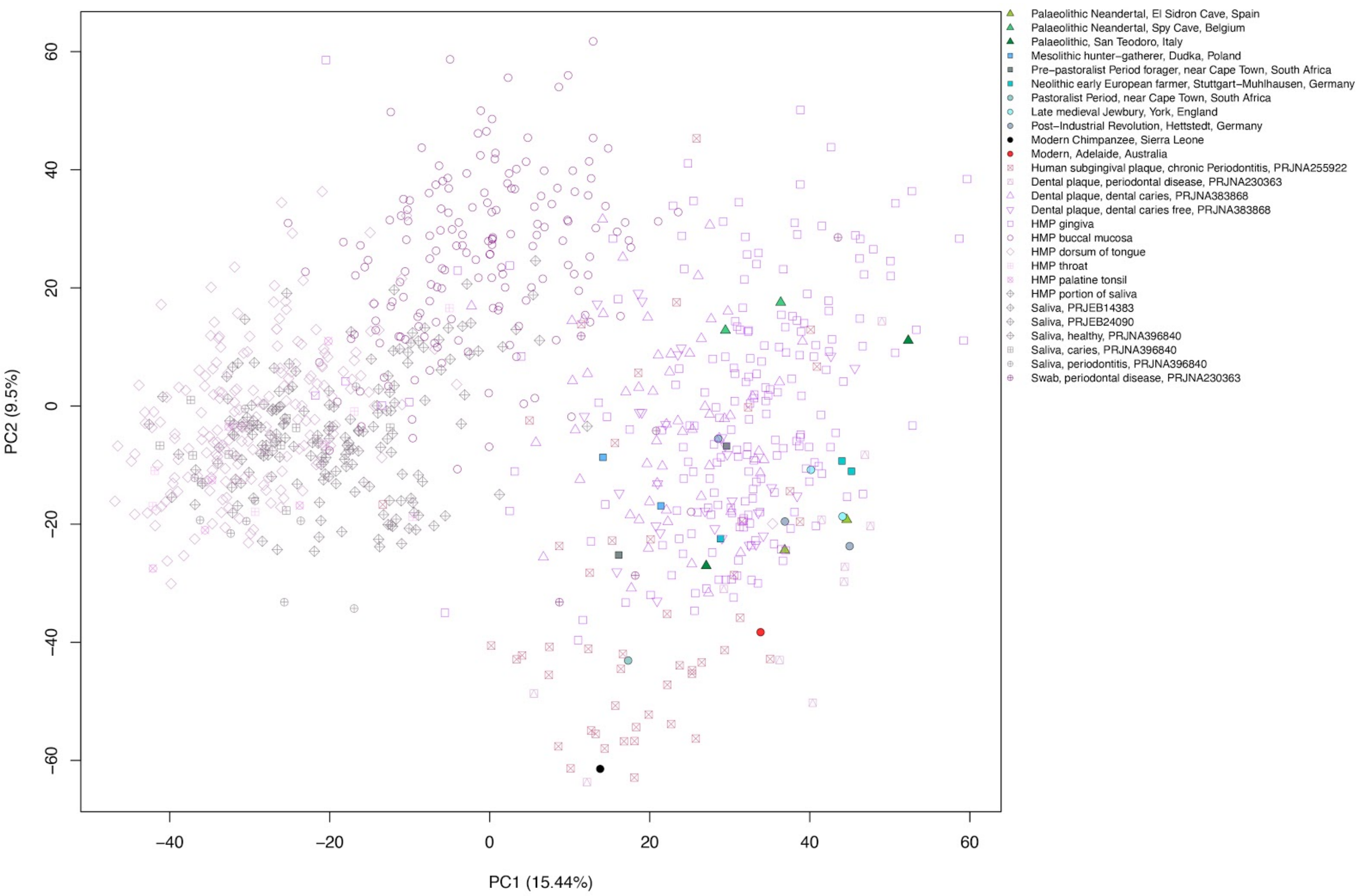


**Fig. S21:** PCA oral samples, clr transformed.

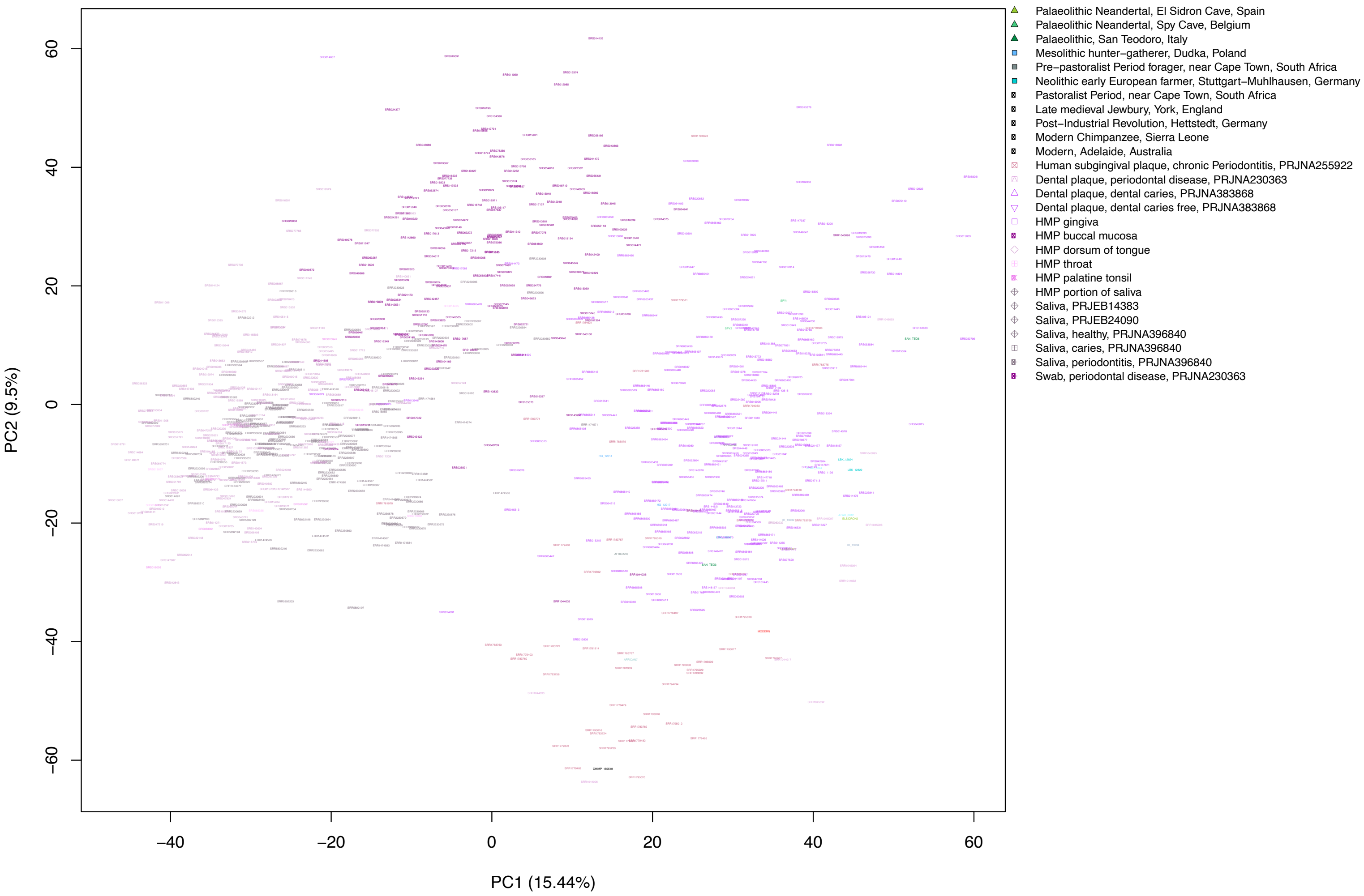


**Fig. S22:** PCA oral samples, including labels.

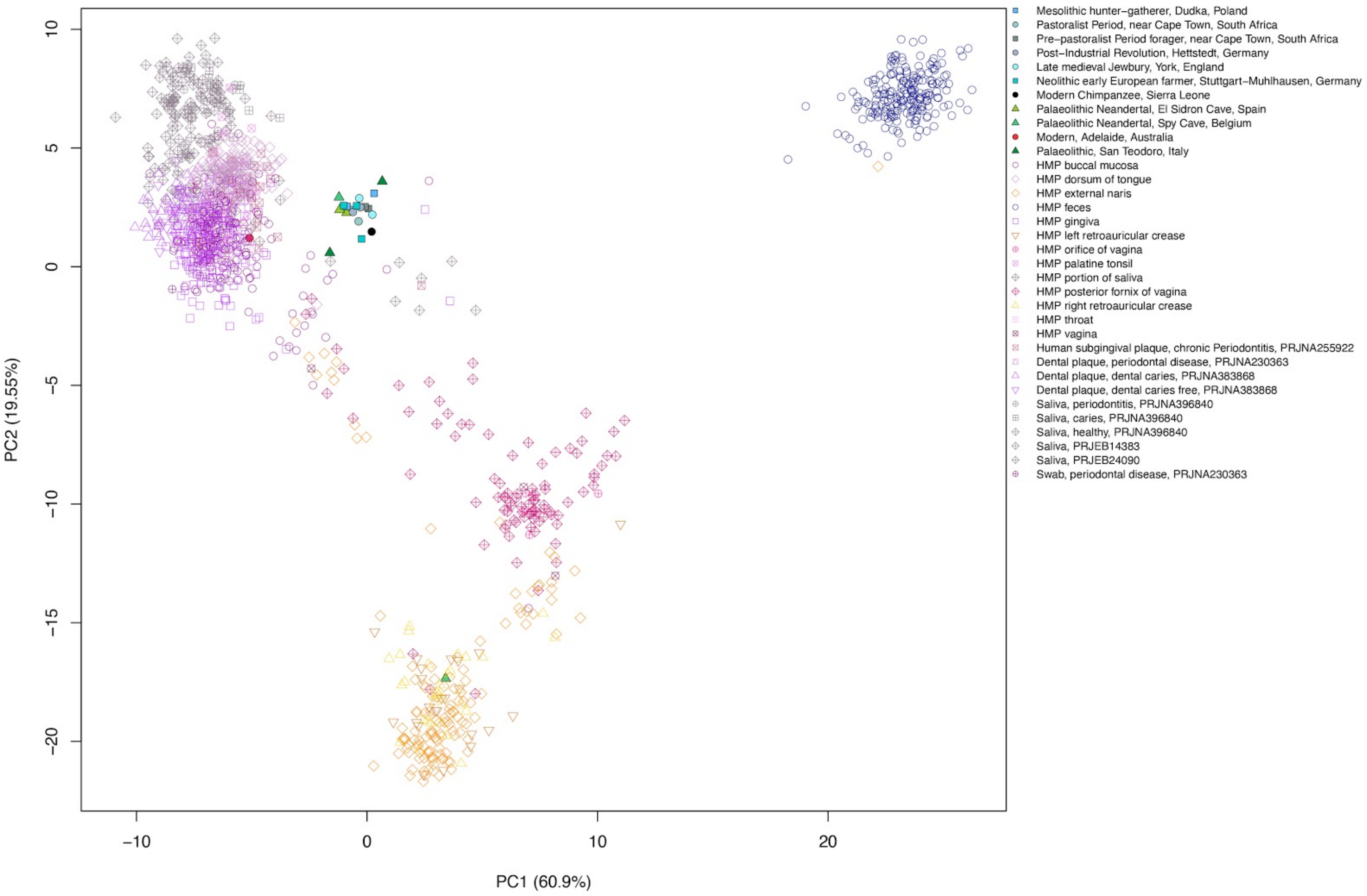


**Fig. S23:** DAPC (DA=15, PC=600), clr-transformed, all samples, grouped by k-means, Table S19.

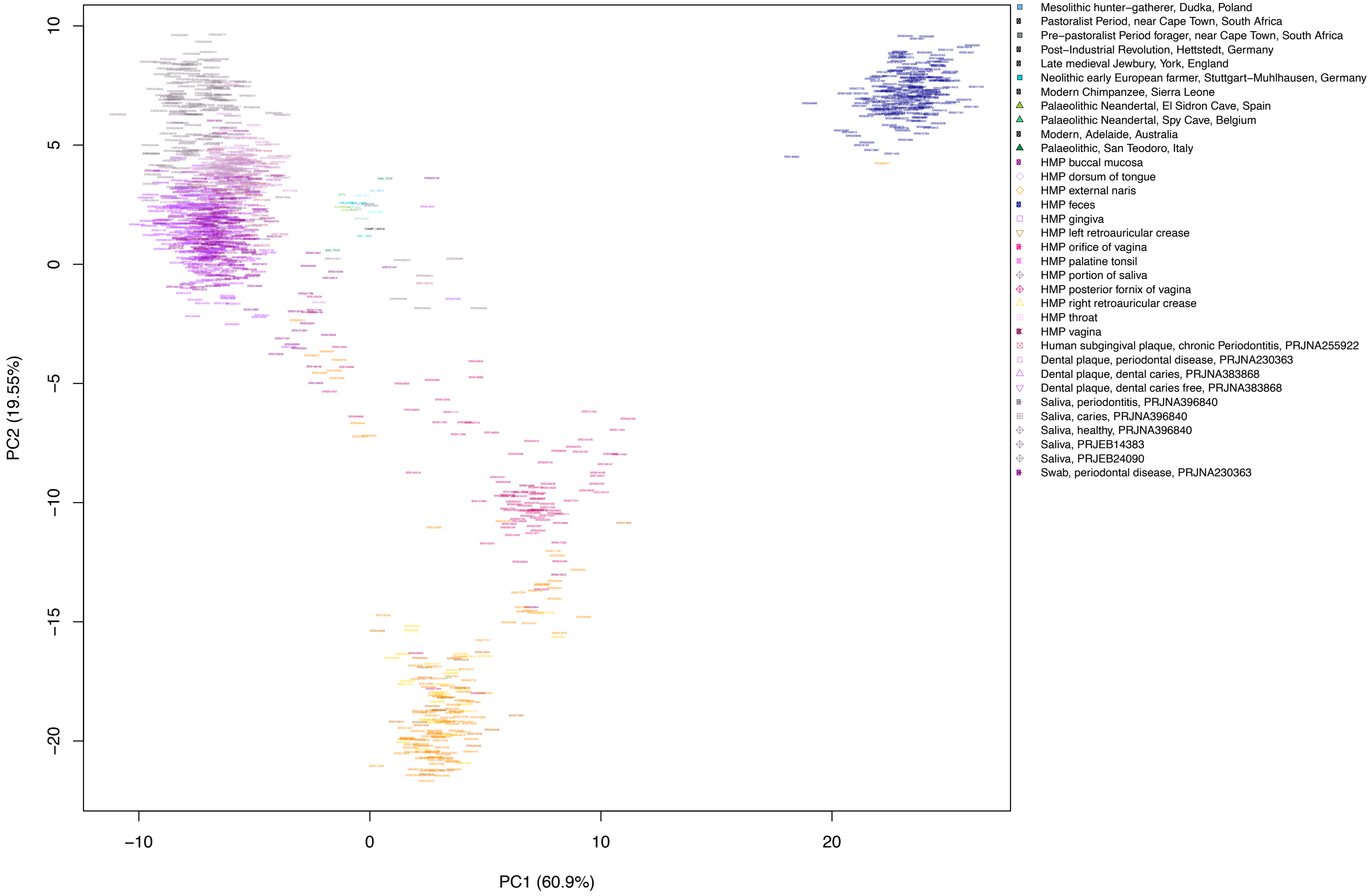


**Fig. S24:** DAPC (DA=15, PC=600), clr-transformed, all samples, grouped by k-means, Table S19. Including labels.

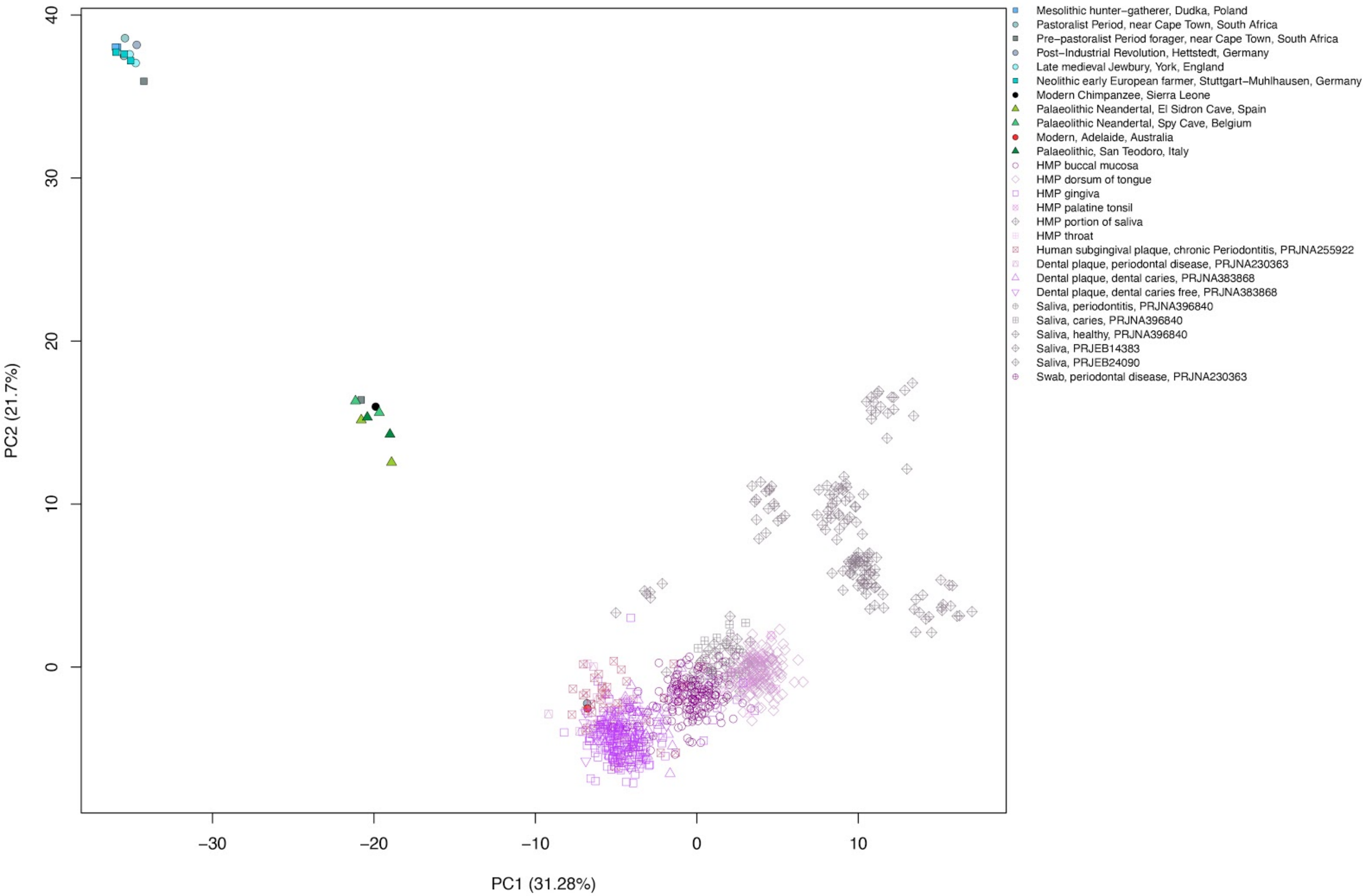


**Fig. S25:** DAPC (DA=15, PC=400), clr-transformed, oral samples, grouped by k-means, Table S20.

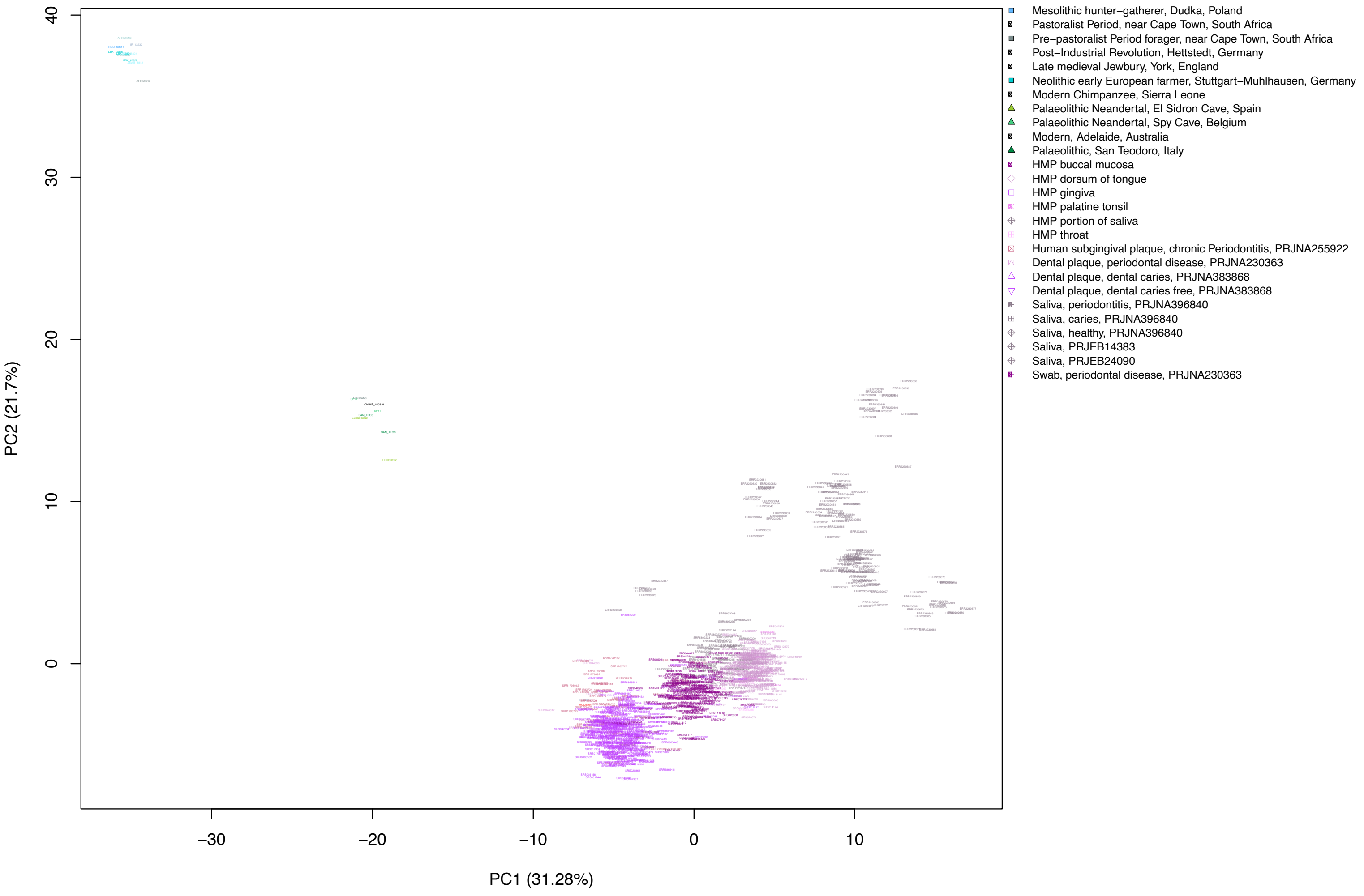


**Fig. S26:** DAPC (DA=15, PC=400), clr-transformed, oral samples, grouped by k-means, Table S20. Including labels.

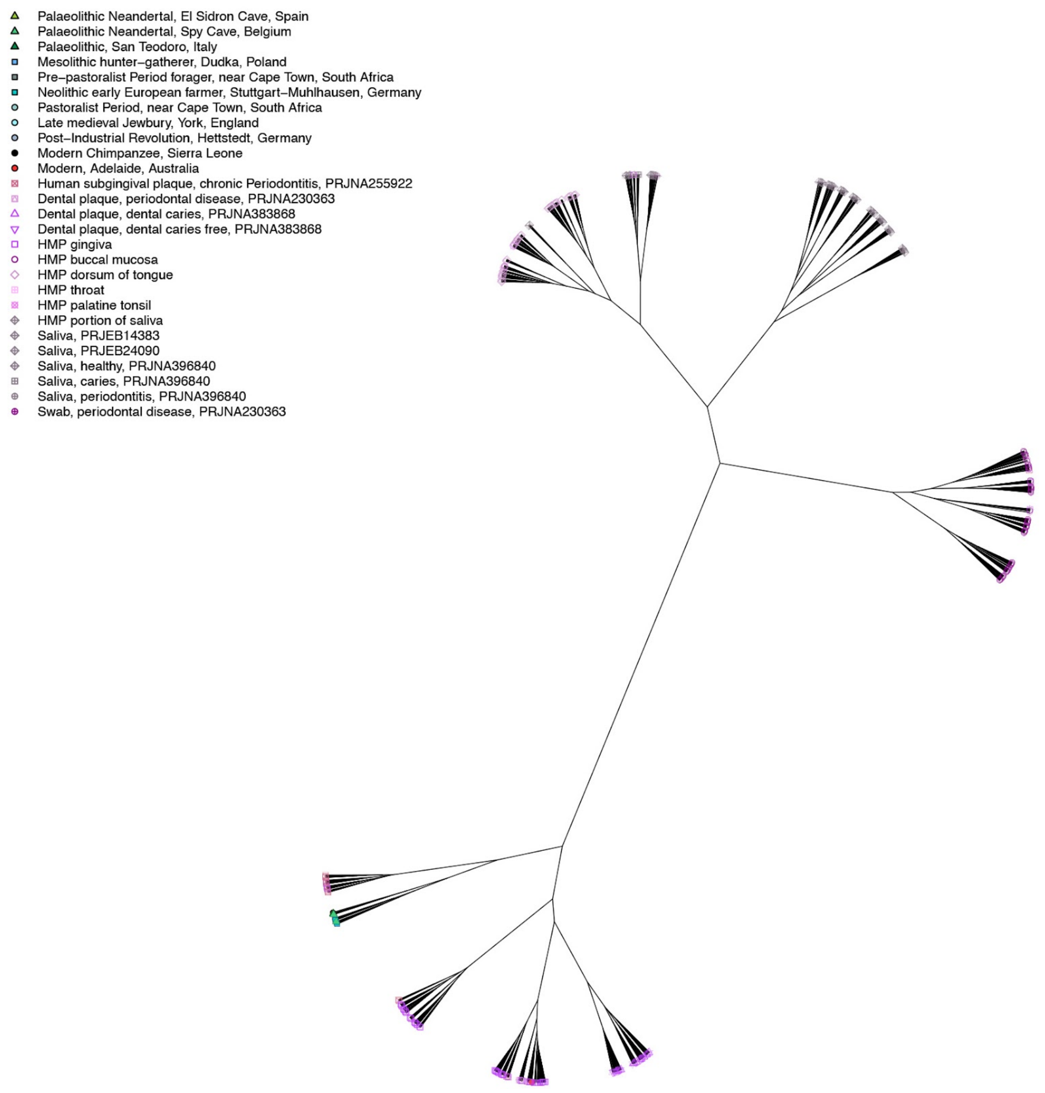


**Fig. S27:** Unrooted dendrogram, clr-transformed, Aitchison distance, oral samples.

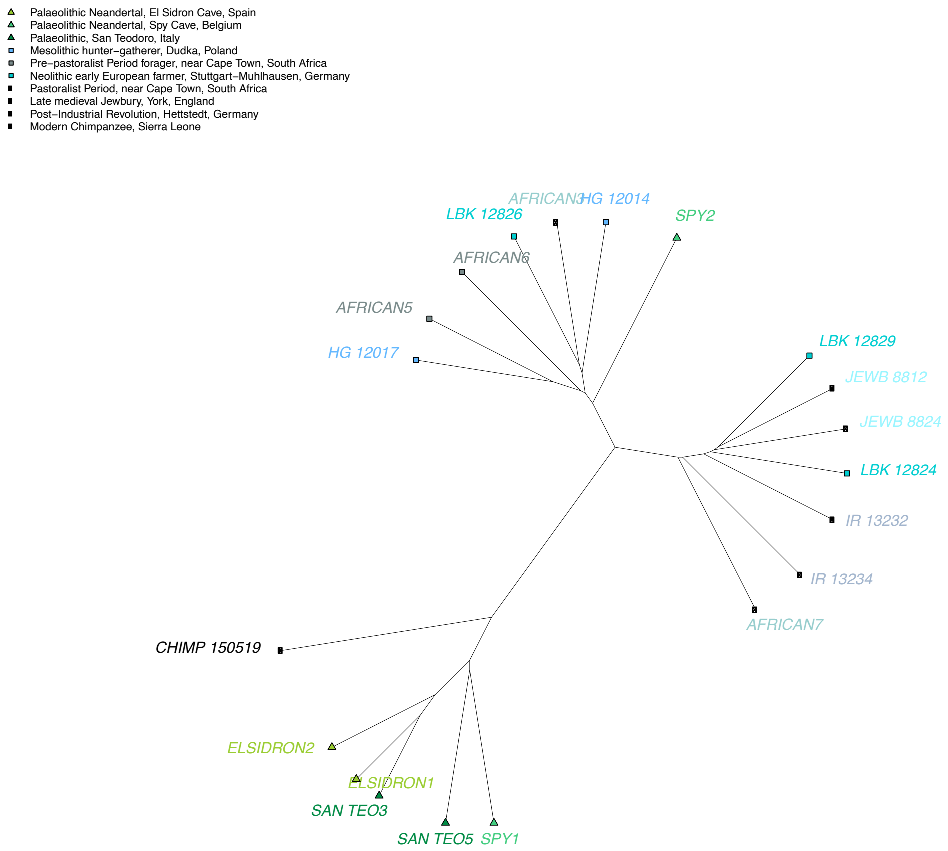


**Fig. S28:** Zoom in on ancient clade in unrooted dendrogram, clr-transformed, Aitchison distance, oral samples.

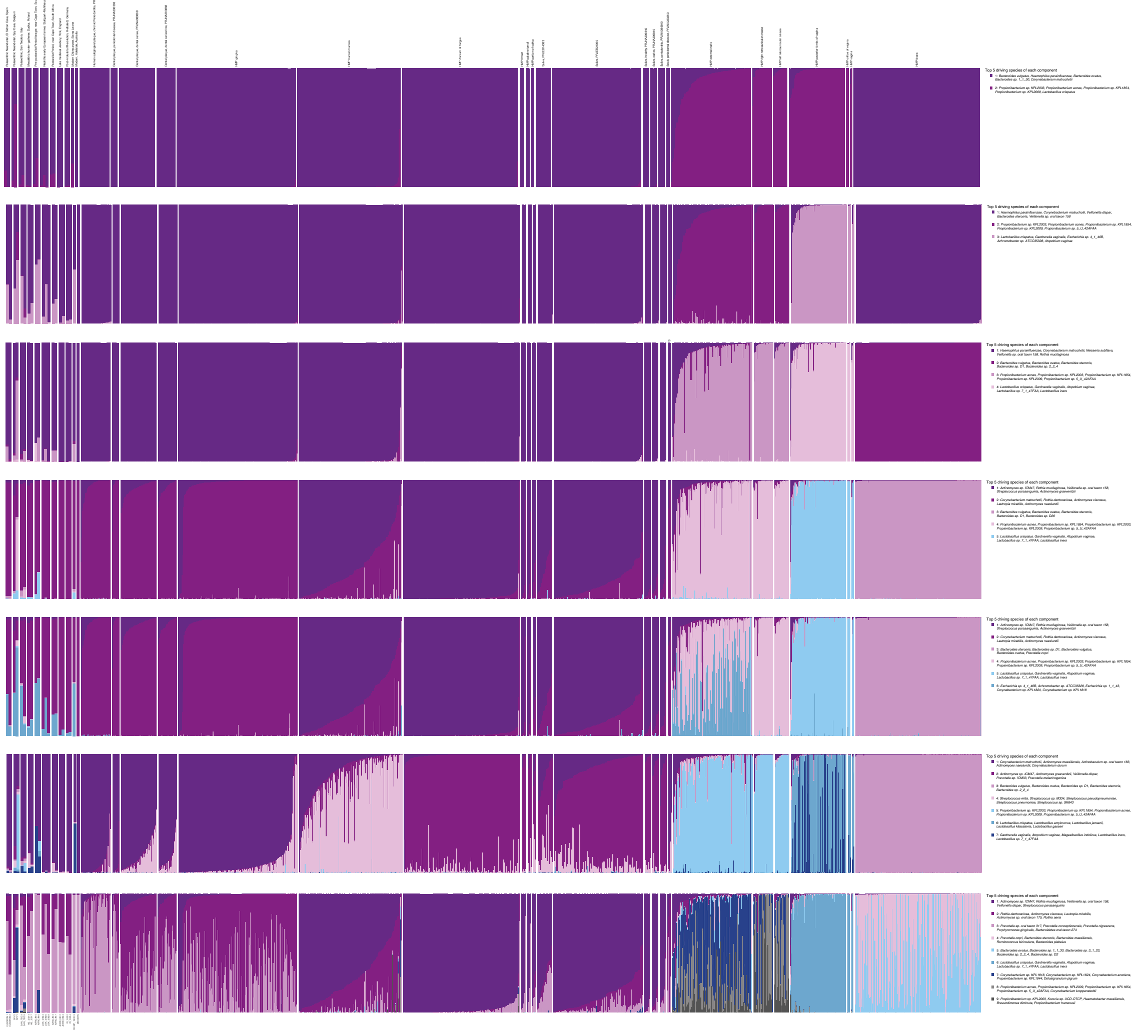

**Fig. S29:** Grade of membership (GoM), all samples.

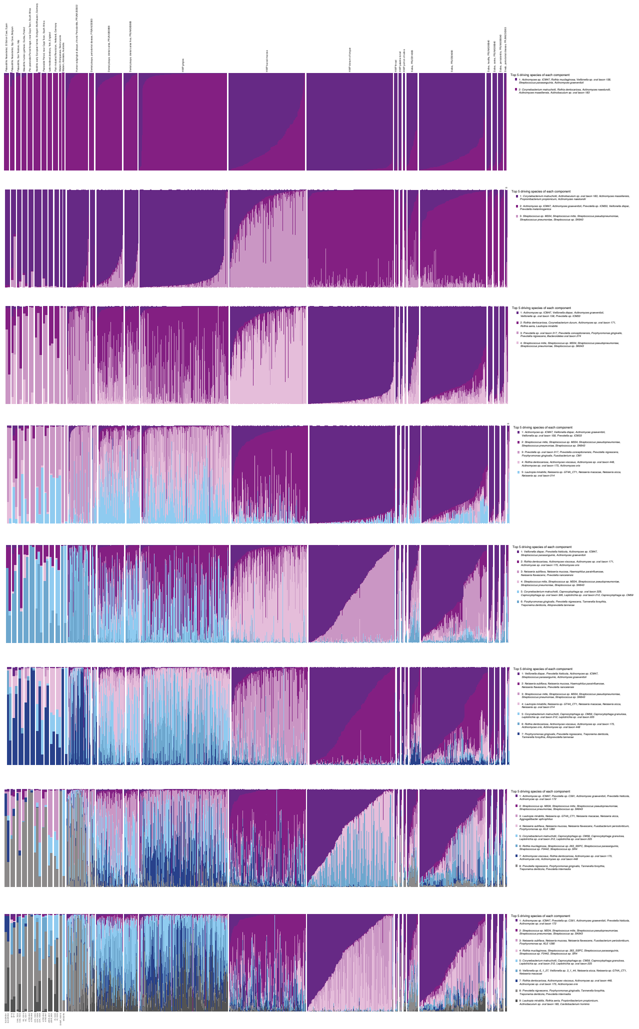

**Fig. S30:** GoM, oral samples.

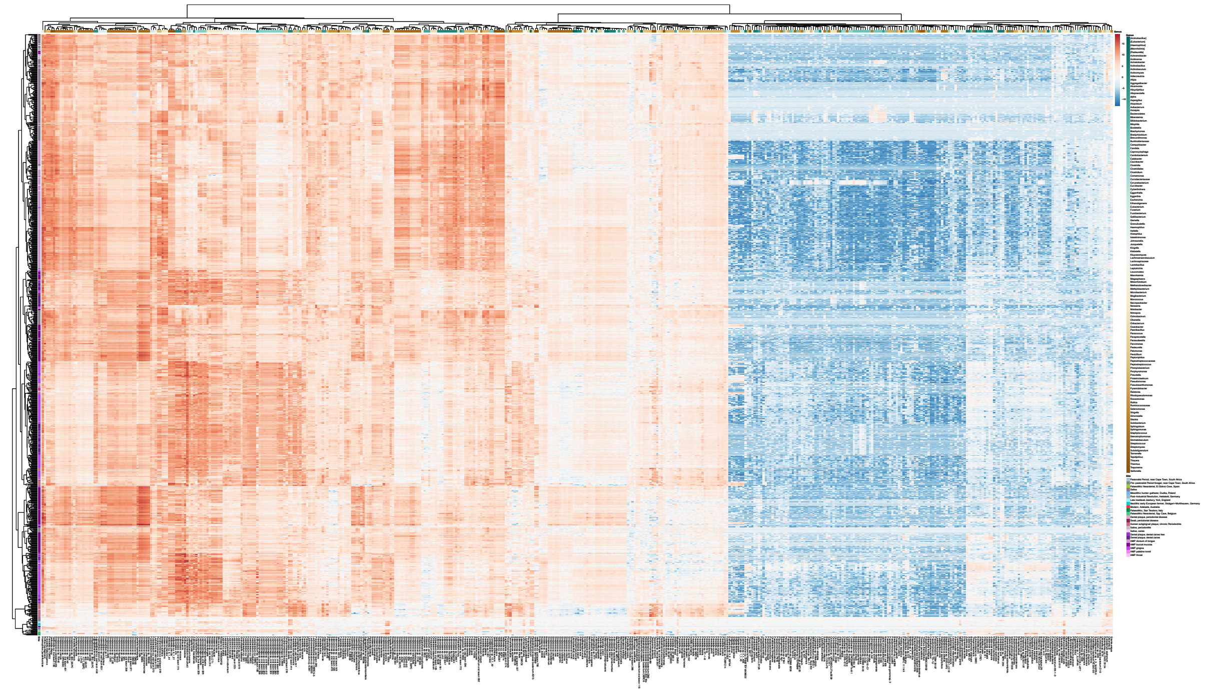

**Fig. S31:** Heatmap of significantly different abundant species (P-value<0.01) between modern oral and ancient calculus samples using ALDEx2. Hierarchical clustering of species on the x-axis and the samples isolation source on the y-axis. Heatmap colored by centred-log transformed abundance estimate. White: the sample has the mean abundance of the species. Blue: the sample abundance of the species is below the mean. Red: the sample abundance of the species is above the mean.

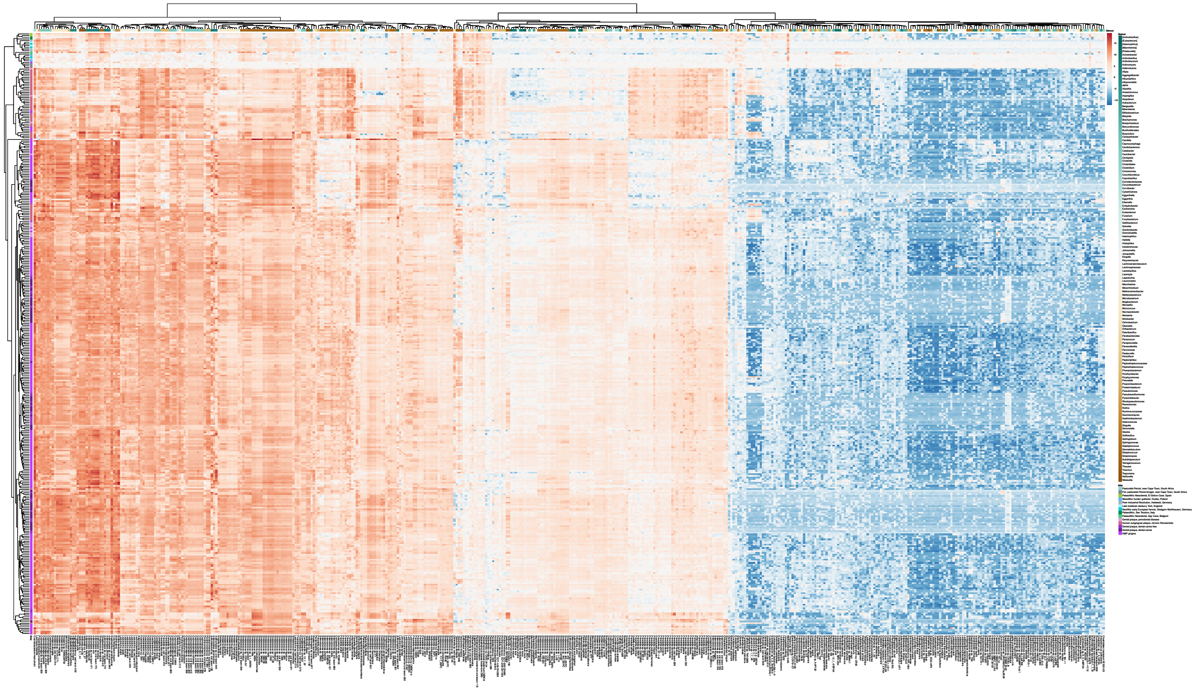

**Fig. S32:** Heatmap of significantly different abundant species (P-value<0.01) between modern gingiva/plaque and ancient calculus samples using ALDEx2. Hierarchical clustering of species on the x-axis and the samples isolation source on the y-axis. Heatmap colored by centred-log transformed abundance estimate. White: the sample has the mean abundance of the species. Blue: the sample abundance of the species is below the mean. Red: the sample abundance of the species is above the mean.

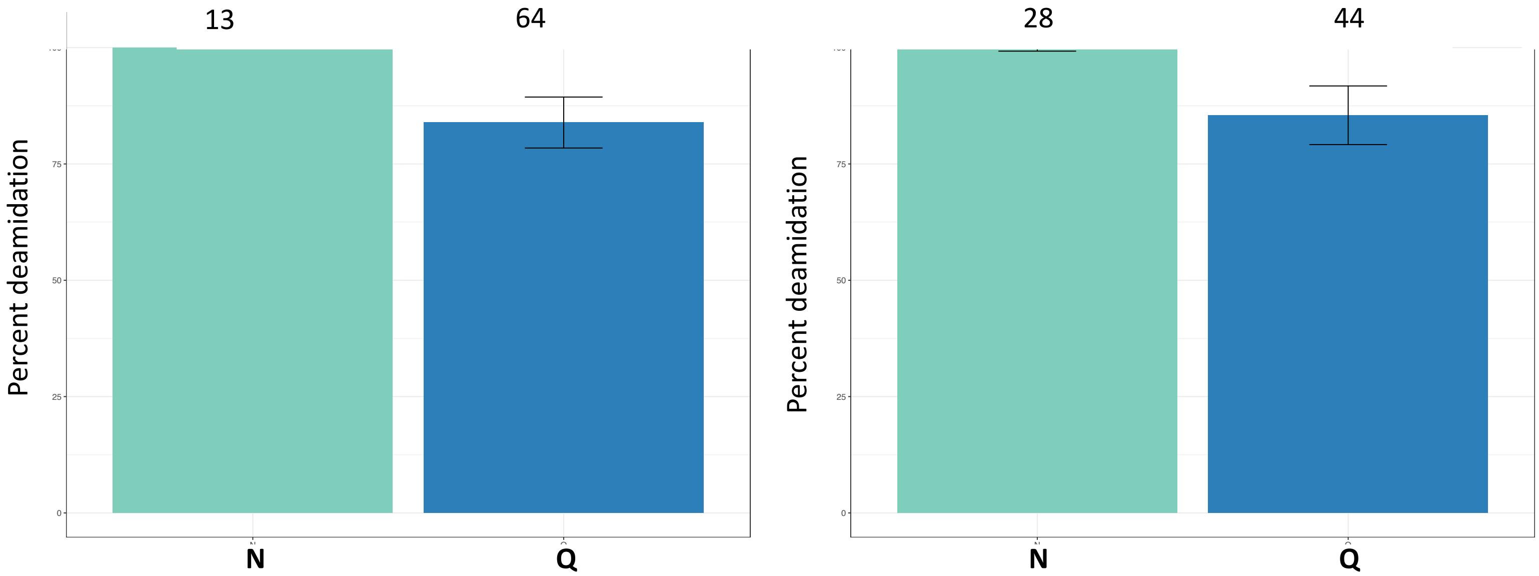


**Fig. S33:** Overall percentage of deamidation for asparagine (N) and glutamine (Q) amino acids for the collagen protein found in San Teodoro 3: a) dental calculus b) petrous bone. Numbers above each bar represent the number of peptides used for the analysis and the error bars represent standard deviation.

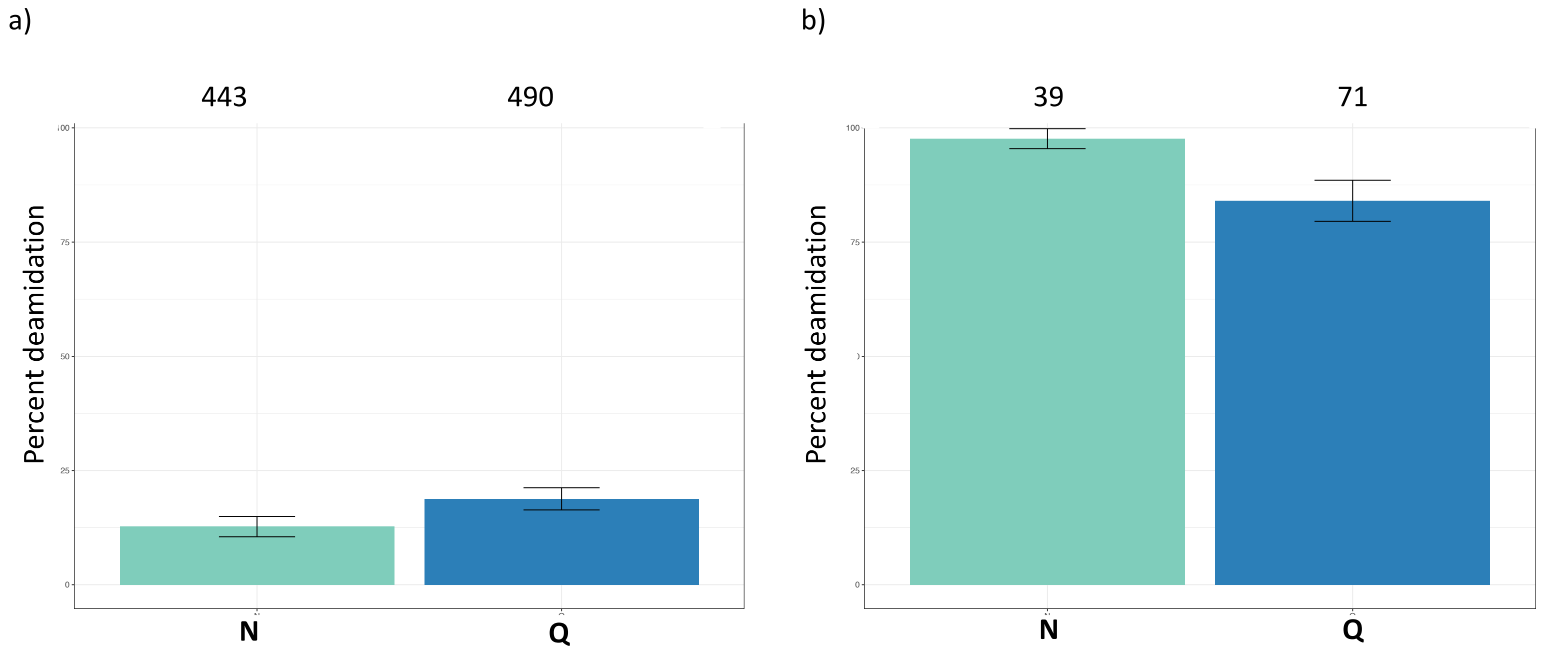


**Fig. S34:** Overall percentage of deamidation for asparagine (N) and glutamine (Q) amino acids for the collagen protein found in San Teodoro 5: a) dental calculus b) petrous bone. Numbers above each bar represent the number of peptides used for the analysis and the error bars represent standard deviation.
